# Context-dependent determinants of CRISPR-Cas9 editing efficiency revealed through cross-species endogenous editing analysis

**DOI:** 10.64898/2026.03.18.712093

**Authors:** Shai Cohen, Shaked Bergman, Adi Yahalom, Alina Wiener, Anat Lavi-Itzkovitz, Anat Maoz, Anna Rice, Dan Dominissini, Dana Furest, David Simanovich Zini, Dongqi Li, Dov Gertz, Ehud Landau, Ester Yeshayahou, Eyal Emmanuel, Gal Avital, Hanin Wattad, Inbar Nevo Yassaf, Itamar Menuhin-Gruman, Karin Brezinger-Dayan, Larisa Rabinovich, Liat Cohen, Mark Bogen, Mark Kapel, Michal Avitzour, Michal J. Besser, Michal Rahimi, Milsee Mol Jaya Prakashan, Mor Cohen Friedman, Moritz Burghardt, Nir Arbel, Ofir Arias, Ofir Yaish, Ofri Kutchinsky, Orna Steinberg Shemer, Rivka Manor, Rivka Ofir, Ruth van-Oss Pinhasi, Shai Izraeli, Shaked Regev, Tal Sherman, Tamara Izhiman, Tom Malul, Vered Chalifa-Caspi, Yehuda Shovman, Yehudit Birger, Amir Sagi, Asaph Aharoni, David Burstein, Eran Eyal, Gilad Kunis, Idan Alyagor, Imri Ben-Israel, Isana Veksler-Lublinsky, Itay Mayrose, Oren Parnas, Oshry Markovich, Sharon Moshkovitz, Shira Corem, Yael Barhum, Yaron Orenstein, Zohar Yakhini, Tamir Tuller

## Abstract

Accurate prediction of CRISPR-Cas9 guide RNA (gRNA) editing efficiency remains limited, particularly outside human systems, where models trained on exogenous human datasets show poor generalization. We analyzed Cas9 efficiency and repair outcomes using novel endogenous editing data from four human cell types, two tomato cell types, and cells from giant river prawn and black soldier fly. While integrating publicly available predictors via ensemble frameworks improved performance, our analysis revealed hundreds of novel features affecting activity. Crucially, dominant features related to sites’ competition for gRNA, and local geometric properties varied across systems, highlighting the strong context dependence of Cas9 efficiency and arguing against a universal model. Interestingly, codon usage bias-based features also emerged as informative predictors, as they are proxies for chromatin accessibility. In contrast, trends in repair outcomes remained conserved. This work provides essential resources for more generalizable CRISPR guide design.

## Introduction

CRISPR-Cas9 has fundamentally changed genome engineering by enabling targeted, programmable DNA modification across diverse organisms (1–7). Mechanistically, Cas9 is directed by a guide RNA (gRNA) to a genomic site immediately upstream of a protospacer-adjacent motif (PAM); a high-fidelity match between gRNA and target generally permits binding and cleavage, after which cellular repair via non-homologous end joining (NHEJ) repair pathway can produce mutations (Fig. 1A). In practice, however, the simple “match-and-cut” rule is insufficient: many gRNAs that satisfy sequence-level rules fail to produce robust edits.

**Figure 1.**
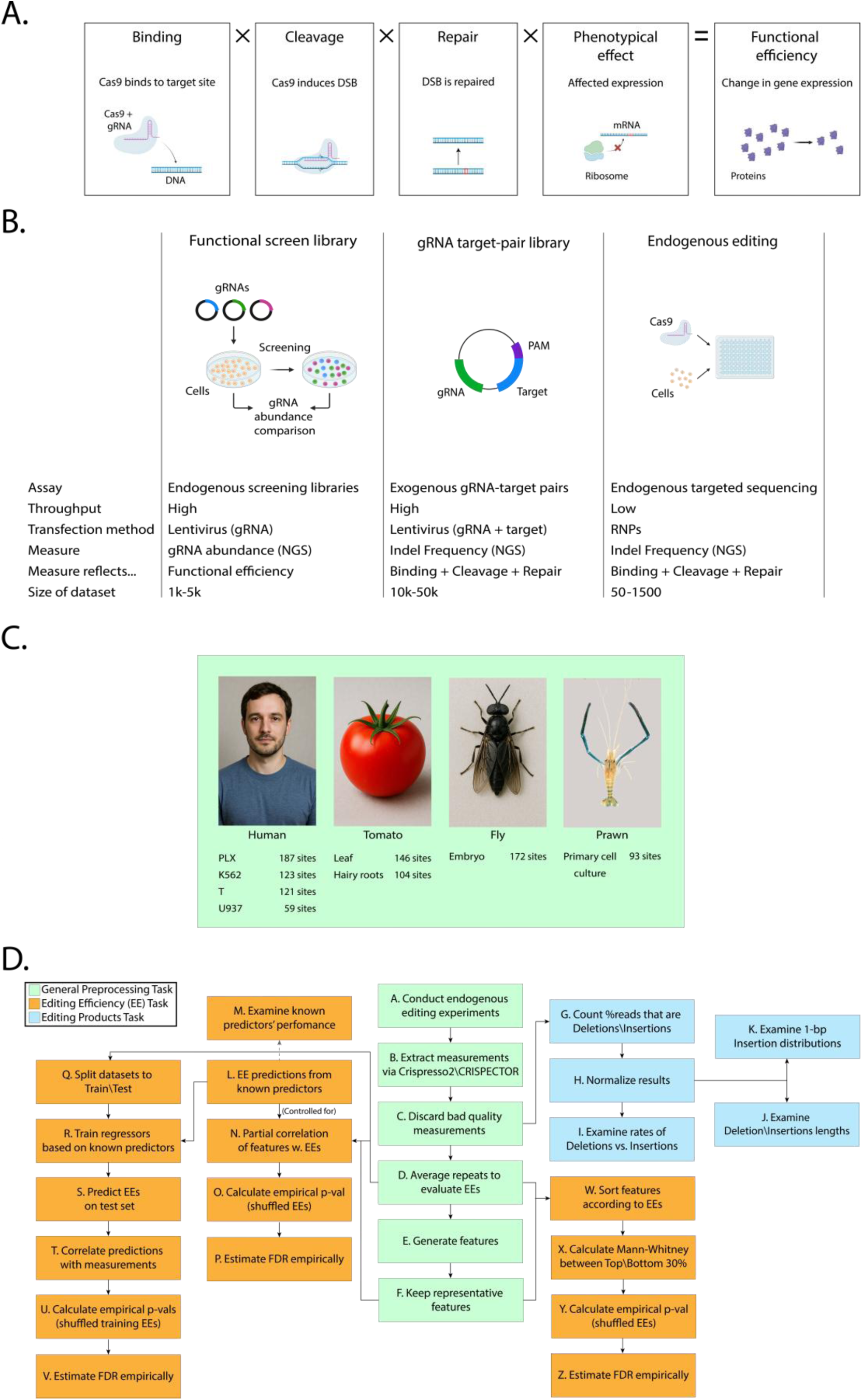
CRISPR experimental methods and details about study. **A.** The problem of predicting the functional effects of gene editing experiments can be decomposed to 4 steps. Combination of the steps 1&2 (Binding & Cleavage) dictate the cutting efficiency. Combining these with Repair i.e. imperfect repair that induces a mutation, defines the editing efficiency. When all 4 steps transpire, they describe the functional efficiency of the sgRNA and usually dictate the experiment’s success. **B.** Three types of CRISPR experiments used to created editing efficiency data. **C.** We conducted Endogenous editing experiments in human, tomato, and black soldier fly. The human cell types were maternal/fetal placenta-derived mesenchymal-like adherent stromal cells (PLX), erythroleukemia cells (K562), T cells, and pro-monocytic histiocytic lymphoma cells (U937). The tomato cell types were leaf protoplasts of M82 tomato from few-weeks old seedlings, and from hairy roots. The fly cell types were multiple embryonic cell types that were not differentiated. The prawn cells were taken from primary cell cultures of 10-14 days old embryos. For details see in methods section “Calculating editing efficiency from experiments outputs”. **D.** General flow chart of the study.

It is important to distinguish three related efficiencies along the pathway from targeting to outcome (Fig. 1A), although they’re often used interchangeably depending on the context: cutting efficiency, the probability that target binding results in a double-strand break; editing efficiency, the probability that a break is repaired into a stable sequence change; and functional efficiency, the probability that an edit produces the intended phenotypic effect. These represent successive steps from binding and cleavage to repair to biological consequence, and failures can occur at any stage.

While there are few computational models predicting gRNA functional efficiency (8), numerous computational models predicting editing efficiency have been developed, including deep-learning approaches trained on large experimental datasets and reported in high-impact journals (9–12). Yet, these models fail to generalize on other data, with correlations frequently dropping to insignificance: within-dataset high accuracy frequently collapses on independent test sets, producing weak often insignificant correlations between predicted and observed efficiencies. A primary reason is bias in the underlying training data, which arises from the experimental pipelines used to collect editing measurements (Fig. 1B). Almost all models are trained on functional screening or gRNA-target pair experiments. Functional screening measures gRNA impact on target gene expression by linking editing to cellular fitness, but this approach is restricted to genes whose disruption affects viability or growth, making it difficult to compare measurements across different loci. In addition, it is dependent on the expression of the payload by the lentivirus carrying the CRISPR-related genes. The gRNA-target pair experiments offer much higher throughput, enabling tens of thousands of measurements, but they assess targets in artificial contexts rather than their native genomic locations, limiting accuracy when predictions are applied in vivo. In contrast, locus-specific endogenous assays provide the most faithful representation of binding, cleavage, and repair within the native genomic context; However, they have significantly lower throughput and require greater experimental efforts and costs. Unfortunately, there are only a few public datasets of this experimental setup, causing a major bottleneck in the field of creating reliable predictors of editing efficiency.

An additional major limitation is taxonomic bias: most publicly available data and models target human systems, yet the CRISPR technology is widely applied in plants, invertebrates and non-model organisms used in basic research, biotechnology and agriculture. As a result, model utility across species is limited and often unknown. Moreover, between different human cell types there is variability in models’ performance with better performance on the cells on which the models were trained.

Understanding the exact properties of repair outcomes can lead to a more appropriate design of gRNAs for various purposes. For example, for knock-out applications it is recommended to select gRNA whose cleavage gives rise to repair products which generate out-of-frame non-functional transcripts (insertions or deletions with length non-divisible by 3). In cleavage designed within CG islands, one might want to prefer gRNAs which give rise to long deletions affecting as many methylation sites as possible.

While software tools for prediction of repair outcome do exist (11, 13–15), they were all trained on human cells only. The nature of repair in other organisms has been overlooked, as well as possible differences in repair outcomes in various human cell types.

To address all the gaps stated above, we performed a cross-laboratory effort that prioritizes endogenous editing assays with RNP (Cas9-gRNA) delivery across multiple human and non-human cell types to reduce protocol-induced biases and better capture binding, cleavage and repair in native chromatin state. We generated data for 4 human cell types, cells in culture of the giant river prawn (*Macrobrachium rosenbergii*), 2 cell types of tomato (*Solanum lycopersicum* cv. M82), and embryos of black soldier fly (*Hermetia illucens*) (Fig 1C). In total, we generated 1297 sites across different organisms and cells, out of which 1005 sites passed post-processing filters (Fig. 1D).

Using these datasets, we engineered diverse, interpretable feature families hypothesized to capture conserved determinants of editing efficiency, including competition/off-target burden, epigenetic and accessibility proxies, thermodynamic and structure descriptors, expression-context metrics, sequence and flanking-sequence features, and metagenomic frequency summaries. Here we present the datasets, features and modeling results, and discuss implications for robust, widely applicable gRNA design.

## Results

### Current editing efficiency predictors do not generalize well across human cell-types or different experimental methods

In recent years many models have been developed for editing efficiency predictions that reported excellent performance on intra-study sets, therefore we chose to analyze the performance of some representative well-known models that were trained on different types of data and relatively well used (9–12). Unfortunately, models trained on data acquired via one experimental method perform poorly when predicting efficiencies on data from other experimental data (Fig. 2A-B). For example, the model trained on data using endogenous editing on T cells (SPROUT), that had spearman of 0.41 on its dataset, performed poorly on datasets from other cell types and high-throughput experimental methods (average Spearman correlation 0.11). As a further testament to this fact, models trained on publicly available datasets with different experimental methods do not perform well on the human datasets that we produced using endogenous editing. For example, DeepCRISPR achieved a 0.96 Spearman correlation on its dataset but on our endogenous human data it achieved an average Spearman correlation of 0.08. In fact, in most cases, these models tend to work well only when tested on the data that was used to train them. Moreover, when tested on non-human datasets their results are also poor.

**Figure 2.**
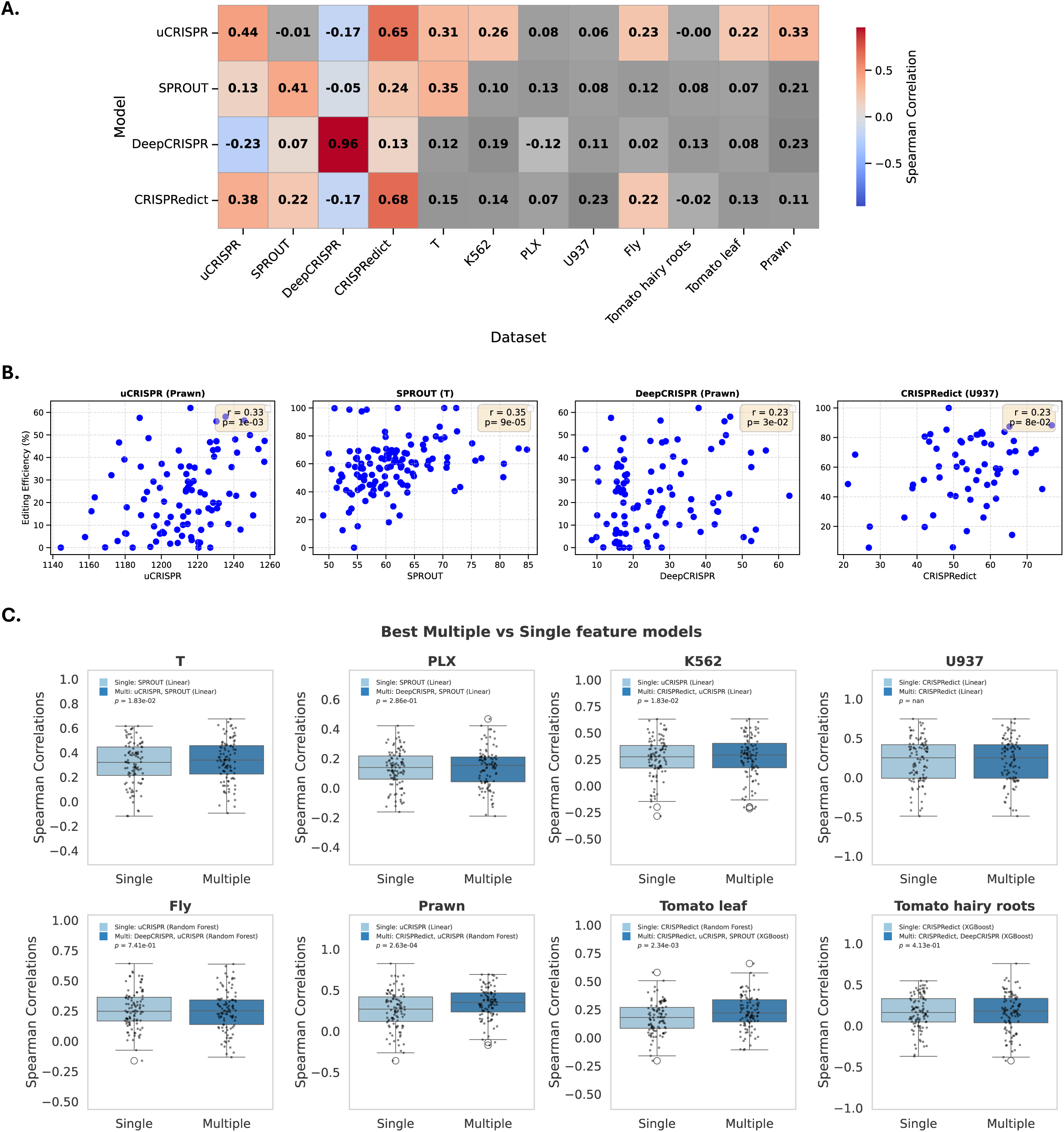
Single and combinations of models’ performances. **A.** Performance of public models on datasets from different cells and experimental methods. Results in gray cells were insignificant. **B.** Models’ prediction and editing efficiency in different cells for cells with maximal correlation. “r” represents spearman correlation. **C.** Results of regression models trained on the data of the current study. The horizontal line in each box represents the mean, and the range are the black lines around each box. Empty dots are outliers. “leave_{X}” means that all other features, but X were chosen.

The models also rarely agree with each other, as the correlations between their predictions are poor (median correlation ranges from 0.18-0.35 over all 8 cells), suggesting that the features learned for every model are overfitted to the training data (Table S1-S8).

### Combination of models usually performs better than single models

Since the publicly available models did not perform well, we tried to improve the prediction based on the data we generated. We trained regression models on our data to predict editing efficiency using the public models’ prediction scores as features (see Methods section for technical details). Our new models outperformed all public models for 7 out of the 8 cells, 4 of which with statistically significant margin (Figure 2C, Table S9).

We verified the results of the regression models by shuffling the editing efficiencies during training (Table S10).

Interestingly, while a linear model provided the best performance in human cells, non-linear models (XGBoost or Random Forest) were superior in fly, tomato, and prawn datasets. This might reflect the fact that the four predictors (CRISPRedict, SPROUT, DeepCRISPR, and uCRISPR) were trained on human data. Consequently, their outputs are well-calibrated and linearly related to true efficiencies in human cells, but exhibit more complex, species-specific deviations in non-human contexts. It seems that non-linear ensemble methods are therefore better able to reconcile these heterogeneous prediction patterns across species.

### Original feature generation

We generated 557 features, most of which are novel, capturing different potential determinants of genome editing efficiency. These features were divided into 5 main categories: (I) Thermodynamics and structure, which account for thermodynamic properties of the DNA target site, based solely on the sequence of the DNA and the gRNA (including its scaffold); (II) Expression features, based on the expression of genes in which the target sites reside; (III) Epigenetics and accessibility features, estimating the physical accessibility of the target site and its 3D density; (IV) Metagenomic features, aggregating data from multiple metagenomics experiments to evaluate the target site’s frequency in CRISPR arrays in nature, providing evolutionary information. (V) Competition features, based on the number of potential off-target sites. We assessed associations between a range of sequence and structure derived features and Cas9 editing efficiency by comparing gRNAs in the top- versus bottom-30% of measured editing efficiency using one-sided Mann-Whitney test (30% was chosen as the threshold as it included enough data in each group to make the comparison, and still makes the two groups different in their editing efficiency range). To obtain robust, data-driven significance estimates that account for feature dependencies and sampling structure, we used a permutation-based framework to derive empirical p-values (technical details in Methods). Applying this procedure revealed system-specific features (Fig. 3) that differentiated between high and low efficiency gRNAs in different cells (Fig. 4, Table S11-S12). Descriptions of all (including non-significant) features and their calculation are presented in Table 1 and in Methods section 2 - “Features generation”.

**Figure 3.**
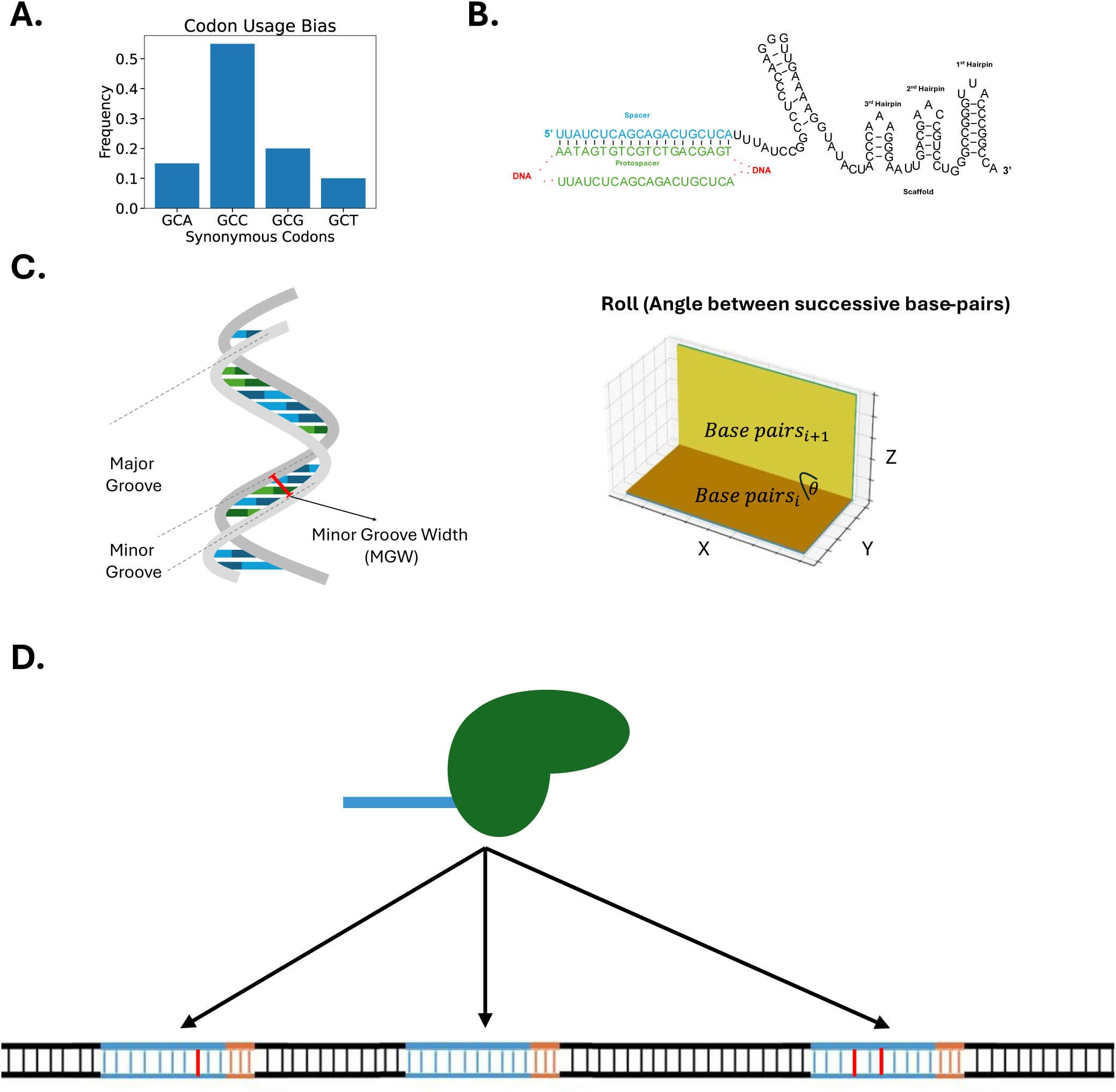
Features we used to predict cutting efficiency in a cell-specific manner. **A.** Codon usage and translational adaptation features calculated from the cell’s own genome, including CAI_avg, chimera_avg, codon_freqs_avg, and aa_freqs_avg, scored across ORF regions overlapping CRISPR target sites**. B.** gRNA thermodynamic and structural features: predicted RNA–DNA melting temperature (gRNADNA), guide RNA free energy including scaffold (guide&scafEne), and predicted scaffold hairpin configurations (Head1, Head2, 1&2&3). **C.** DNA shape features at target sites, including minor groove width (MGW) and Roll, derived from DNAshape predictions. **D.** Competition-based features quantifying nearby partially matching sequences at different genomic scales, with or without NGG PAM motifs.

**Figure 4.**
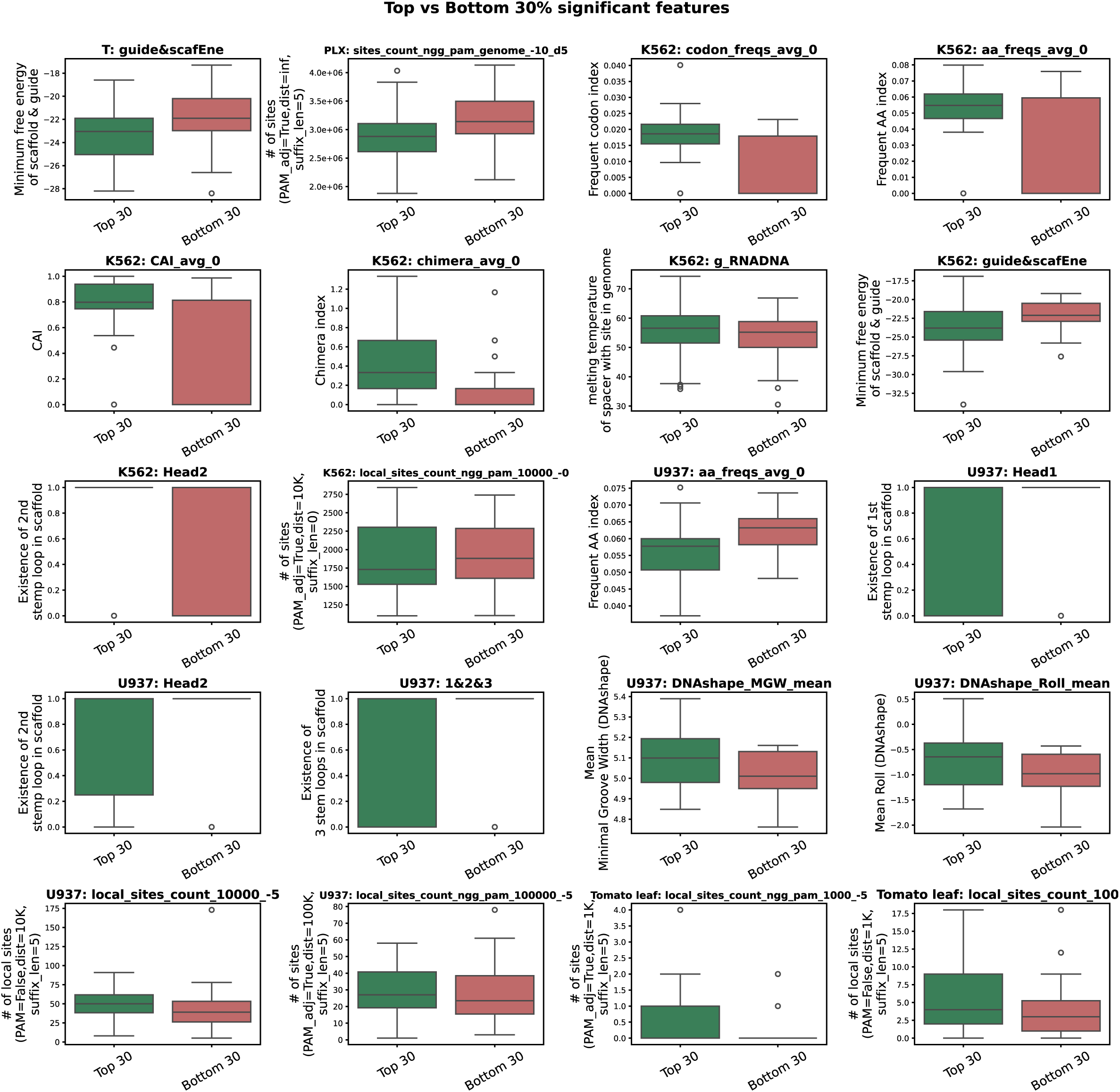
Features with significant differences in high efficiency gRNAs compared to low efficiency gRNAs in various cell types. For each cell & feature, we sorted the feature values according to the editing efficiency and compared the top 30% of the feature values with the bottom 30% using a one-sided Mann-Whitney test. The box plots are of the Q2-Q3 percentiles of the values in the top\bottom 30%. Horizonal lines in each box is the mean, and the range are the black lines around each box. Empty dots are outliers. All results are significant with an p-value of 0.0008.

**Table 1:** Short description of all 557 features.

**Short description of all 557 features - Table 1:**
| Feature | Details |
| --- | --- |
| enthalpy_mean / enthalpy_mean_ext /<br>enthalpy_mean_up/down_ext / enthalpy_x | Average or position-specific nearest-neighbor DNA enthalpy values reflecting folding strength around the target site. |
| DNAshape_{MGW/HeIT/ProT/Roll}_mean / _mean_ext /<br>_mean_up/down_ext / _x | Local or extended average or position-specific DNA shape features (minor groove width, helical twist, propeller twist, roll) predicted from pentamers (18). |
| Scaffold stem features (“Head1”, “Head2”, “Head3”,<br>“1&2&3”) | Binary indicators of whether the gRNA scaffold forms specific predicted RNA secondary-structure canonical stem loops using ViennaRNA’s RNAFold (19, 20). |
| g_RNADNA\DNADNA | Gibbs free energy ($\Delta G$, in kcal/mol at the specified temperature) of duplex formation for the guide spacer hybridizing to its complementary strand as an RNA:DNA hybrid (g_RNADNA) or as a |
|  | DNA:DNA duplex (g_DNADNA), reflecting the thermodynamic stability of each pairing. |
| **guideEne** | Minimum free energy of the gRNA alone using ViennaRNA's RNAFold (19, 20). |
| **guide&scafEne** | Minimum free energy of the gRNA plus scaffold complex using ViennaRNA's RNAFold (19, 20). |
| **isBasePairs** | Binary indicator of whether gRNA nucleotides are predicted to base-pair with scaffold nucleotides using ViennaRNA's RNAFold (19, 20). |
| **DNA_fold** | Predicted folding free energy of the genomic site $\pm 10$ nt computed from MATLAB's oligoprop thermodynamic tables. |
| **aa_freqs** | Geometric mean of genomic amino-acid frequencies for codons overlapping the window around the target site. |
| **codon_freqs** | Geometric mean of genomic codon |
|  | frequencies for codons overlapping the window around the target site. |
| **CAI** | Codon adaptation index<br>- measuring how frequently overlapping codons appear relative to optimal synonymous codons (21). |
| **chimeraARS** | Compatibility score depicting how many long sub-strings in highly expressed genes are present in the local window (22). |
| **Methylation-motif score** | Maximum match score between local sequence windows and empirical methylation motifs from (23). |
| **k-mer quartile features (3-mers & 5-mers × 5 spacer sets)** | Fraction of gRNA k-mers falling into each frequency quartile derived from metagenomics CRISPR spacer datasets. |
| **k-mer log-probability features** | Log-likelihood of observing the gRNA's k-mers in each metagenomics CRISPR spacer set. |
| **Reduced-alphabet (GC/AT) k-mer features** | Same quartile and log-probability k-mer metrics computed after |
|  | collapsing nucleotides into GC vs AT categories. |
| **Local Composition Complexity (LCC)** | KL-divergence–based measure of distinctiveness of nucleotide composition in sliding windows around the target (24). |
| **sites_count_ngg_pam_{ref_seq}_{suffix_length}_{d}** | Number of genome/ORF/gene sites matching the target’s PAM-adjacent suffix (5–20 nt) with ≤d mismatches (0–5) followed by NGG. |
| **local_sites_count_{ngg_pam}_{max_distance}_{suffix_length}** | Number of nearby competing sites (0–1,000,000 nt away) matching suffixes of various lengths (0–5 nt or PAM-only), optionally requiring NGG PAM. |

Briefly, the features that captured cell-specific determinants of cutting efficiency (Fig. 3-4), included features based on codon usage bias, translational adaptation, RNA thermodynamics, DNA structural properties, and potential local competition effects (Fig. 3). Codon usage bias was quantified using the codon adaptation index (CAI_avg), which measures the preferential usage of synonymous codons relative to the most frequent codon in the genome. In parallel, the Chimera score (chimera_avg) estimates how well an ORF is adapted to the host translational machinery by assessing the presence of long substrings commonly found in native coding regions. We additionally included codon_freqs_avg and aa_freqs_avg to quantify codon and amino acid frequencies across the proteome. All codon-related measures were calculated using reference sets derived from each cell’s own genome and were scored across ORF regions overlapping the CRISPR target sites. Because higher codon adaptation is typically associated with elevated gene expression, these features may indirectly reflect chromatin accessibility (16) and transcriptional activity.

To characterize gRNA-intrinsic properties, we computed the predicted melting temperature between the RNA spacer and its complementary genomic DNA target (gRNADNA), reflecting hybridization stability. We also calculated the free energy of the full guide RNA including its scaffold (guide&scafEne) as a measure of overall structural stability. Predicted scaffold hairpin configurations (Head1, Head2, and 1&2&3) were included to account for variation in canonical secondary structure formation.

DNA geometric features were derived from DNAshape predictions. Minor groove width (MGW) was calculated as the mean across all 19 overlapping pentamers within each target site, capturing local groove geometry that may influence Cas9–DNA recognition. Similarly, the Roll parameter, which reflects the inclination between adjacent base pairs and local DNA flexibility, was included because such structural properties can affect Cas9 binding and cleavage efficiency.

Finally, we defined competition-based features to quantify nearby genomic sequences partially matching the CRISPR target site. These features counted local matches at multiple genomic scales (1 kb–100 kb), with or without NGG PAM motifs, and included genome-wide counts allowing mismatches within the PAM-adjacent suffix. Such partially matching sites could modulate editing efficiency either by facilitating local Cas9 diffusion and transient binding near the true target or by sequestering Cas9 away from it. Despite experimental evidence supporting these competitive effects (17), they have not, to the best of our knowledge, been incorporated into predictive models of gRNA efficiency.

Our analysis revealed a striking heterogeneity in the determinants of Cas9 activity across biological systems, suggesting that predictive features are highly sensitive to cellular context. In K562 cells, high-efficiency guides were primarily characterized by coding-context features, with significant enrichment observed for average codon frequency (codon_freqs_avg), amino-acid frequency (aa_freqs_avg), and Codon Adaptation Index (21) (CAI_avg).

Conversely, local genomic context in K562 cells appeared to exert an inhibitory effect; the density of PAM-adjacent sites (local_sites_count_ngg_pam) was significantly higher among low-efficiency guides. This negative effect is consistent with a “decoy” model, where a high density of near-cognate sites locally sequesters Cas9 or sterically hinders effective cleavage at the target locus. In contrast, U937 cells displayed a distinct feature profile dominated by structural properties and an inverted relationship with target density. DNA shape attributes, specifically mean Minor Groove Width (DNAshape_MGW_mean) and Roll (DNAshape_Roll_mean), were significant positive predictors of efficiency (see Figure 4). Notably, unlike in K562, local site density features (e.g., local_sites_count_10000_-5) in U937 cells were positively correlated with high editing efficiency. This positive association was observed also in the plant system, where tomato leaf protoplasts similarly exhibited higher local site counts among high-efficiency guides. Thermodynamic stability metrics related to guide folding and scaffold stems (e.g., guide&scafEne, Head1/3) failed to show a universal unidirectional effect, functioning as positive predictors in some datasets and negative in others, further reinforcing the idiosyncrasy of these systems. The opposing effects of local site density - deleterious in K562 but beneficial in U937 and tomato - argue against a single, universal mechanism for Cas9 target search and cleavage.

### Partial correlation analysis reveals features with predictive power orthogonal to previous models

To determine whether the system-specific associations identified in the previous section represent a signal not already explained by existing traditional CRISPR efficiency predictors, we performed a complementary analysis using partial Spearman correlations between each feature and measured editing efficiency while controlling for four public models: SPROUT, uCRISPR, CRISPRedict, and DeepCRISPR. For each feature, we computed the Spearman correlation between the residuals of the feature and the residuals of editing efficiency after regressing out these four predictors, thereby isolating the unique contribution of the feature (see technical details in the Methods section and all the results in Table S13-S14). The analysis revealed that many (8 out of 20) of the features identified in the Mann-Whitney framework retain their association with editing efficiency even after conditioning on four state-of-the-art predictive models. Specifically, in K562 cells, all coding-context features showed strong negative partial correlations: codon_freqs_avg_0 (r = −0.46, p = 0.0001), aa_freqs_avg_0 (r = −0.34, p = 0.0001), CAI_avg_0 (r = −0.42, p = 0.0001), and chimera_avg_0 (r = −0.46, p = 0.0001). These results indicate that the robust associations observed in Fig. 4 are not captured by SPROUT, uCRISPR, CRISPRedict, or DeepCRISPR, and instead reflect a model-independent signal. In PLX, isBasePairs showed a positive partial correlation (r = 0.09, p = 0.0018).

Local-site-count metrics (i.e. competition-based features) also remain significant in several different systems. local_sites_count_ngg_pam_1000_-0 exhibited a negative partial correlation (r = −0.18, p = 0.0079), and sites_count_ngg_pam_genome_-10_d5 showed a positive association (r = 0.28, p = 0.0001). In Tomato leaf protoplasts, local_sites_count_ngg_pam_1000_-5 also showed a significant negative partial correlation (r = −0.15, p = 0.0097). These results align with the context-dependent directions observed in Fig. 4, reinforcing the notion that PAM-adjacent similarity has system-specific biological interpretations.

Guide-folding and structural features likewise displayed significant residual effects. In T cells, Head3 showed a positive partial correlation (r = 0.15, p = 0.0031), while local_sites_count_ngg_pam_1000_-0 (r = −0.28, p = 0.0012) and local_sites_count_ngg_pam_10000_-0 (r = −0.31, p = 0.0007) showed strong negative correlations. In U937, multiple folding descriptors were significant, including g_DNADNA (r = 0.34, p = 0.0056), Head1 (r = 0.33, p = 0.0055), and 1&2&3 (r = 0.32, p = 0.0066). In Tomato hairy roots, DNAshape_HelT_mean showed a strong negative association (r = −0.27, p = 0.0028).

Finally, in Tomato leaf protoplasts, coding-context features again showed consistent positive partial correlations: codon_freqs_avg_0 (r = 0.23, p = 0.0025), aa_freqs_avg_0 (r = 0.21, p = 0.0086), CAI_avg_0 (r = 0.25, p = 0.0013), and chimera_avg_0 (r = 0.20, p = 0.0023), matching the directionality seen in Fig. 4.

Overall, these results demonstrate that multiple sequence, structural, and genomic-context features retain significant associations with editing efficiency after controlling for four leading prediction models, indicating that the system-specific determinants observed in Fig. 4 reflect a novel biological signal beyond what current CRISPR prediction frameworks capture.

### Repair profiles are universal in various cellular systems

For each cell other than tomato hairy roots, we obtained short-read amplicon sequencing data of the edited region and used Crispresso2 to analyze NHEJ repair outcomes in the vicinity of the Cas9 cut sites. We then analyzed the list of detected products and event frequencies as reported by the software. Similarly, we obtained long-read amplicon sequencing data from tomato hairy root cells and analyzed it using CRISPECTOR (25), since it is more adequate for that type of data.

Specifically, we were interested in analysis of indel lengths and indel patterns generated by NHEJ repair mechanisms.

We first examined the robustness of amplicon sequencing for repair outcome analysis. Replicates show highly similar repair nature in each case (Fig. 5A), suggesting that the pattern is not random and reflects real characteristic cellular events which govern the repair.

**Figure 5.**
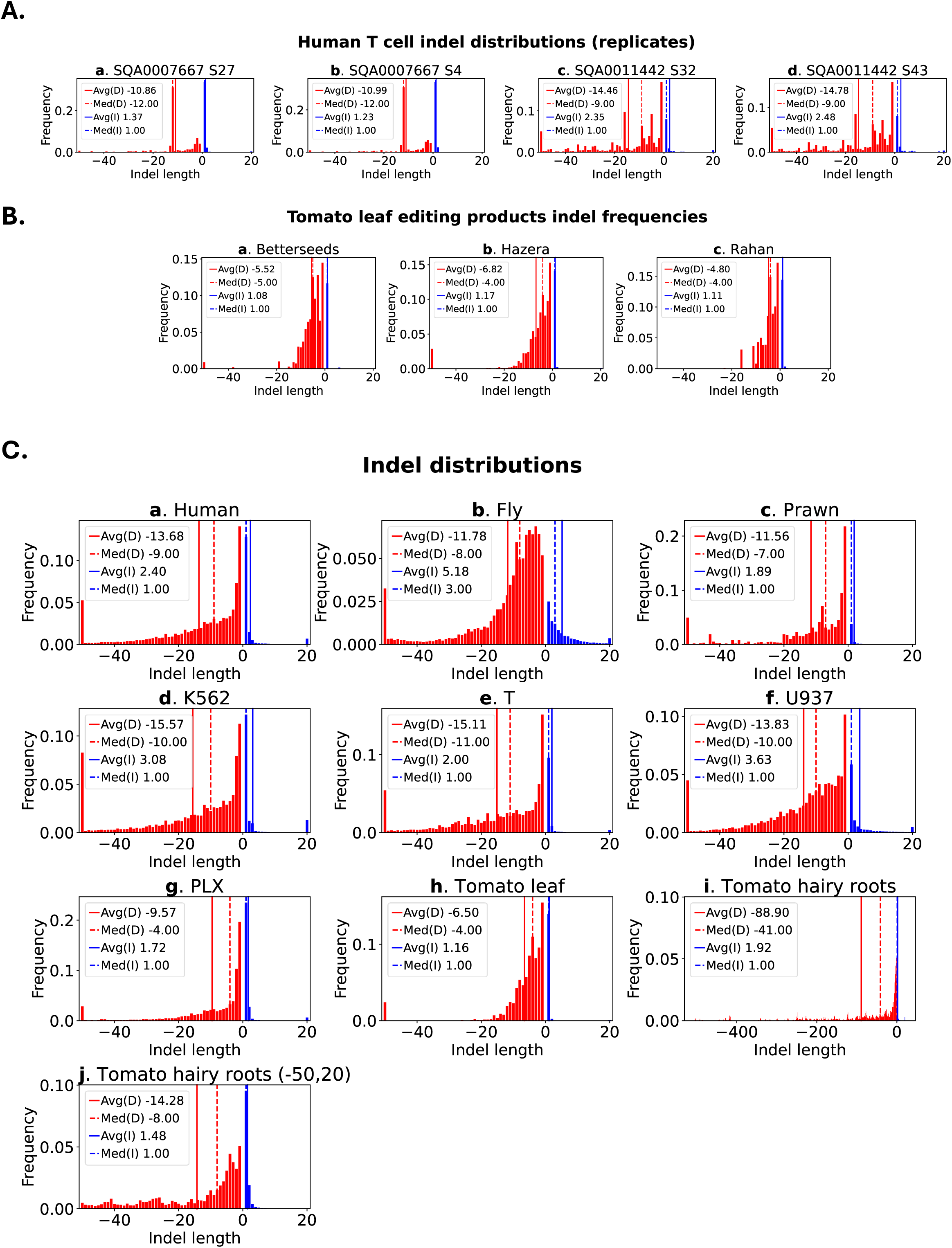
Indel analysis across different cells, organisms, and types of sequencing. In all graphs “Avg” stands for the average, “Med” stands for the median, “D”/”I” stands for deletions/insertions respectively. Positive indel lengths indicate insertions and negative lengths indicate deletions. **A.** Reproducibility of NHEJ editing repair. Shown are repair-product lengths frequencies of biological replicates for two gRNAs (a, b first pair, and c, d second pair). The replicates show highly similar repair nature in both cases. **B.** Repair products frequencies for Tomato leaf protoplasts conducted by different laboratories of biotechnology Ag companies. The product frequencies for all experiments of each company are summed according to the indel lengths. **C.** Repair products frequencies for various types of organisms and cells. The product frequencies for all experiments of each cell type are summed according to the indel length. Events with lengths outside the axis limits are summed in the external bins of the histograms (except for in subfigure k). Panel ‘a’ includes data for all human cells combined (U937, T, K562, PLX) combined. Panel ‘i’ has a different range since it its data was sequenced differently (see “1.2 Methodology for creating figure 5” for more details). Panel ‘j’ is a zoomed-in version of panel ‘i’ for deletions of length 1-50 and insertions between 1-20. p-values for the claim that there are more deletions than insertions were less than 1e-260 for all cells (including “Human”). The spearman correlations between lengths of deletions and insertions ranged from -0.61- -0.95 with significant p-values (See methods section for details). The claim that 1-based insertions are more frequent than other insertions, and that they are usually a copy of the upstream nucleotide are statistically significant and these analyses can found in detail in the methods section.

Fig. 5B shows the similarity in the indel length distribution between 3 experiments for the same cell type (Tomato leaf protoplasts in this case) when conducted by different laboratories - BetterSeeds, Hazera and Rahan. Similar profiles suggest that the repair patterns are not very sensitive to variabilities in the experimental setting and protocols.

We then analyzed the distribution of events over many sites (59-187 sites) for various cellular systems. Fig. 5C (a) shows the indel length distribution for various human cells. The patterns are clearly different between different cells, but in all we observe more deletions than insertions, more short deletions than long deletions and among insertions a clear advantage for 1-bp insertions (see technical details in the Methods section), consistent with previous observations of repair by NHEJ (26–32). The correlations and p-values can be seen in Table S15.

Differences appear mainly in the spectrum of deletion events. A broad spectrum of indel lengths with high frequencies of events (both deletions and insertions) is observed in fly embryos (Fig. 5C (b)). Of all cells investigated this is the only system for which 1-bp deletion was not the most frequent type of deletion. A major difference in the distribution of lengths occurs also with the same organism, as exemplified for human cells here.

### In all cell types and organisms, the predominant single-base insertion significantly corresponds to a duplication of the upstream nucleotide to the cleavage site

Across most cell types studied in this work, the predominant single-base insertion (1-bp insertion) significantly corresponds to a duplication of the upstream nucleotide to the cleavage site (Fig. 6 and technical details in Methods). This pattern is remarkably consistent, suggesting a universal mechanism rather than organism-specific variation, potentially reflecting the staggered-end nature of Cas9 cleavage and subsequent templated fill-in during end joining, as previously observed in mammalian cells (33–36).

**Figure 6.**
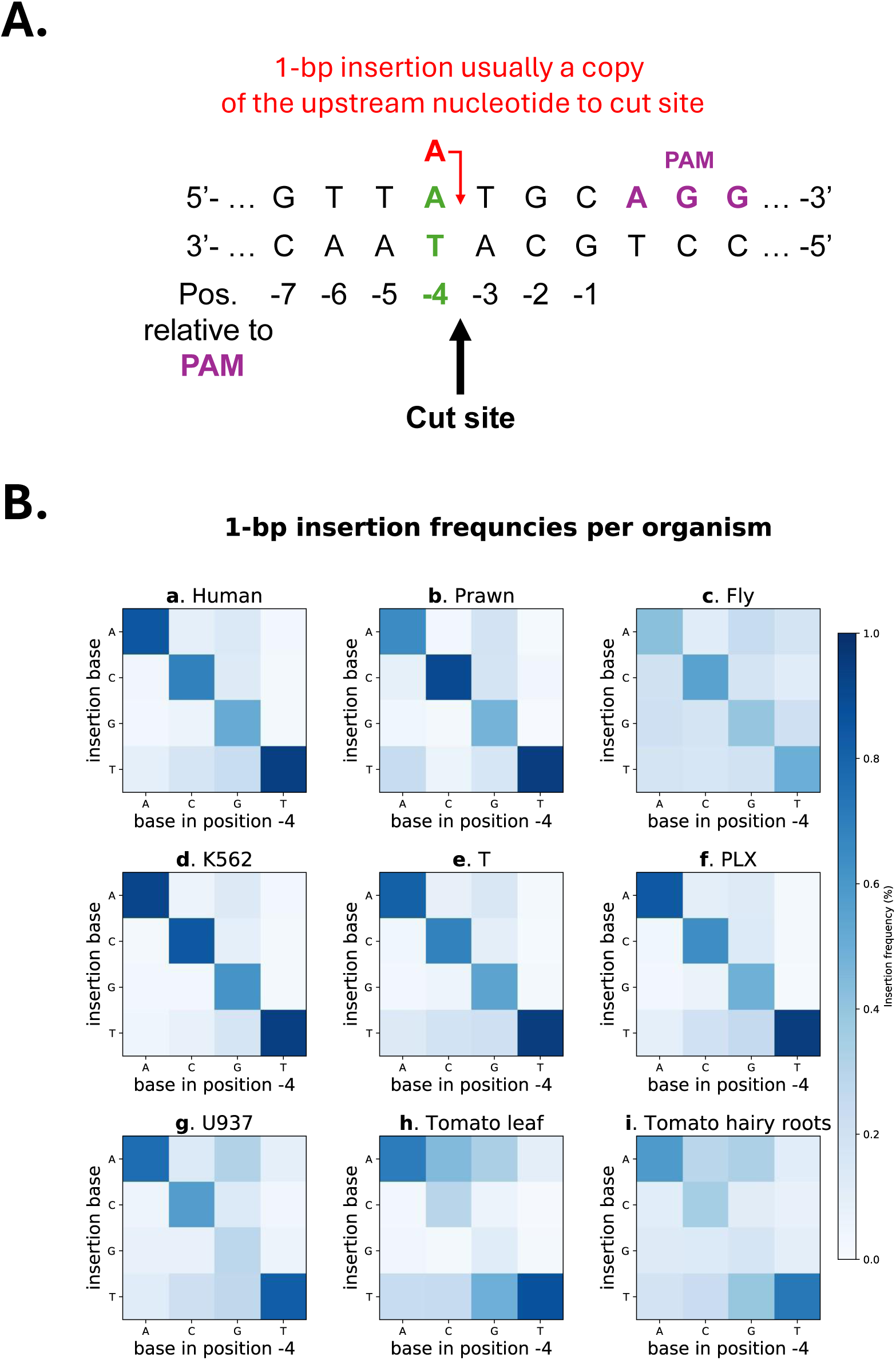
1-bp insertion base identity as a function of the identity of the base upstream to the cut site (position -4) in various organisms. **A.** Illustration of 1-bp insertion that is a copy of the nucleotide that is directly upstream to the cut site, i.e. in position -4 relative to the PAM. **B.** Shown are frequencies, in multiple experiments of all 1-bp insertion products, for the insertion of each type of base (the y-axis) given the type of base immediately upstream to the intended cleavage site (x-axis). The p-values (after FDR correction) for the claim that the inserted nucleotide in 1-bp insertions is under selection to be a copy of the nucleotide upstream to the cut-site for all cells for all nucleotides was smaller than 0.0011 except for both tomato cell types for inserted nucleotide G which was smaller than 0.0032.

Similarly, in human cells examined here (Fig. 6B (‘a’, ‘d’-‘g’)), most 1-bp insertions are copies of the preceding base. These observations indicate that, regardless of organism or cell type, the identity of a 1-bp insertion is strongly determined by its upstream nucleotide. Interestingly this tendency is stronger when there is a pyrimidine base in the upstream position, although it the signal is weaker when G appears in this position for tomato and U937 human cells.

## Discussion

To enable more reliable genome engineering across biological systems, we systematically dissected determinants of Cas9 editing efficiency and repair outcomes using endogenous editing data generated in four human cell types, two tomato cell types, and cell systems derived from giant river prawn and black soldier fly. Because most publicly available CRISPR datasets originate from human functional screens or gRNA-target pair libraries, current predictors are trained predominantly on indirect or exogenous measurements. Consistent with this bias, leading prediction tools showed weak generalization to our datasets. However, we employed ensemble strategies integrating outputs from multiple models improved performance in most systems (7 out of 8), indicating that existing frameworks capture complementary but incomplete signals. Linear ensembles performed best in human cells, whereas non-linear approaches provided greater gains in non-human systems, highlighting species-specific deviations that are not well represented in human-trained models.

Across 557 candidate features spanning sequence, structural, energetic and genomic properties, editing efficiency emerged as a systems-level phenotype shaped by both intrinsic DNA features and local chromatin context. Coding-context features, including codon usage bias (CUB) metrics and amino-acid frequency, were associated with Cas9 editing efficiency in human cells. CUB-derived features were significantly associated with higher editing efficiency in one human cell type. We selected these features because: A. They serve as indirect proxies for chromatin accessibility and local regulatory state (16, 37–42), which are key determinants of Cas9 target recognition and cleavage (32, 43, 44); B. Direct measurements of chromatin accessibility (e.g., ATAC-seq, DNase-seq) are not available for most of the cell types and organisms we examined, particularly in non-model systems. In contrast, amino-acid frequency showed significant associations in two human cell types but with opposite effect directions, suggesting that these features capture regulatory signals interpreted differently across cellular contexts. DNA-shape descriptors such as minor groove width and roll also correlated with efficiency, consistent with models in which local DNA deformability modulates Cas9 binding and R-loop formation (45). PAM-adjacent site density and competition-based features exhibited system-dependent effects, including opposite correlations across cell types, potentially reflecting differences in chromatin organization that influence accessibility of competing genomic sites, although experimental biases may also contribute.

(16, 37–42)(32, 43, 44)(45)Partial correlation analyses controlling for leading public predictors demonstrated that many (8 our of 20) system-specific features retained independent explanatory power, underscoring that substantial, cell-type-dependent biological variation remains unmodeled in current frameworks. Together, these findings support the development of next-generation predictive tools that integrate transcriptional context, DNA structural properties, genomic repetitiveness and guide-level features, rather than relying solely on sequence-derived heuristics.

Across all organisms examined, NHEJ-mediated repair outcomes exhibited strikingly conserved architectural features. Deletions consistently outnumbered insertions, short deletions predominated over longer events, and 1-bp insertions overwhelmingly dominated the insertion spectrum. In most systems, the most frequent 1-bp insertion corresponded to duplication of the nucleotide immediately upstream of the cut site, consistent with minimal end processing and templated fill-in during NHEJ. Similar global biases have been reported in mammals, plants and fish (27–31, 46) but largely from functional screens or exogenous reporter systems. To our knowledge, such conserved repair signatures have not previously been demonstrated using high-quality endogenous editing data across this breadth of taxa, nor in emerging non-model systems such as giant river prawn and black soldier fly. The reproducibility of these patterns across biological replicates, laboratories and sequencing strategies indicates that they reflect intrinsic properties of NHEJ rather than methodological artefact. Although longer deletion events were present in the hairy roots data, they are likely underrepresented in systems analyzed exclusively by short-read sequencing. Consistent with this interpretation, long-read analyses in other organisms have revealed similar extended deletion profiles (47, 48), suggesting that such events are a general feature of NHEJ repair and would likely be observed in additional cell types if assessed using long-read approaches.

From a biotechnology perspective, these conserved repair biases provide actionable guidance for gRNA design across species. The predictable dominance of short deletions and templated 1-bp insertions increases confidence in frameshift induction strategies and informs editing of regulatory or CpG-rich loci. More broadly, our results demonstrate that endogenous editing datasets provide a more faithful representation of in vivo Cas9 behavior than high-throughput screening approaches alone, which may introduce context-specific biases or overlook regulatory features that influence editing outcomes in native chromatin environments. Systematic comparison of functional screens, gRNA - target pair assays and endogenous editing measurements for the same guides within the same cellular background would help quantify the extent to which experimental modality shapes apparent editing efficiency and model performance.

Looking forward, predictive frameworks may benefit from explicitly modeling target-site competition in a chromatin-aware manner. Current approaches typically estimate competition based on sequence similarity alone; however, off-target sites embedded in closed chromatin are less likely to recruit Cas9 and therefore may not meaningfully compete with the intended target. Incorporating chromatin accessibility into competition metrics could refine estimates of effective target availability and improve guide selection accuracy in vivo.

As genome editing expands into agriculture, aquaculture and industrial biotechnology, systematic generation and public release of endogenous datasets in diverse organisms will be essential for building predictive models that generalize beyond human systems and enable rational, cross-species genome engineering.

## Methods

### 1.1 Calculating editing efficiency from experiments outputs

#### 1.1.1 Short-sequencing

For experiments in Human, Fly, Prawn, and Tomato leaf, we analyzed the amplicon sequence reads with the CRISPResso2 (49) algorithm. Experiments that had less than 10K mapped reads or whose percentage of mapped reads was less than 90% were discarded. Furthermore, CRISPResso2 ran with an optimal window of 8 surrounding the cut site (50) and while disregarding substitutions (since dealing with NHEJ products). Editing efficiencies in replicate experiments were averaged for that over the replicates.

#### 1.1.2 Long-sequencing (Tomato hairy roots)

##### 1.1.2.1 CRISPR editing analysis using CRISPECTOR

Sequencing data were processed and analyzed using CRISPECTOR (25, 51) to quantify CRISPR editing outcomes. raw input data consisted of: (i) a FASTQ file containing mock (wild-type) reads, (ii) a pooled FASTQ file containing reads from all edited samples with replicate-specific barcodes, and (iii) an annotation table specifying amplicon identifiers and associated guide RNA sequences (Table S16).

##### 1.1.2.2 Demultiplexing and read normalization

Edited reads were demultiplexed by barcode to generate replicate-specific FASTQ files. To reduce read-depth imbalance between edited and mock samples, the mock FASTQ file was randomly subsampled to one-sixth of its original size and used consistently as the control input across all analyses.

##### 1.1.2.3 Guide-specific amplicon filtering

Amplicons targeted by a single gRNA were processed directly using their corresponding guide sequences as provided in the annotation table.

For amplicons targeted by two gRNAs at distinct genomic locations, reads were separated by target site. As CRISPECTOR does not support simultaneous analysis of multiple cut sites within a single amplicon, reads corresponding to each target location were filtered into separate datasets, labeled T1 and T2, ensuring that each analysis included only a single cleavage site per amplicon.

##### 1.1.2.4 CRISPECTOR execution and batching

Filtered datasets (i.e. T1 & T2) were further divided into batches to manage input size and computational constraints. In total, eight independent CRISPECTOR runs were performed for dual-guide amplicons (separately for T1 and T2 targets), in addition to a single run for amplicons targeted by a single gRNA. All runs used the same subsampled mock control. Summary table for all runs is provided in (Table S17).

CRISPECTOR output reports from all runs were collected and merged for downstream analysis of on- and off-target editing events.

##### 1.1.2.5 Handling repeats

Editing efficiencies in replicate experiments were averaged for that over the replicates.

### 1.2 Methodology for creating figure 5

All plots except ‘i’ & ‘j’ were created by taking from each sample the percentage of reads that had a deletion\insertion and adding them up across all samples and then normalizing them by their total joined sum. The data for plots ‘i’ & ‘j’ came from a different experimental setup (where there were sometimes 2 sites in the same amplicon) which required additional pre-processing (described in the next paragraph). If an experiment was repeated, only a single arbitrary replicate was taken for the short-sequencing data (everything except tomato hairy roots since for the long sequencing data i.e. tomato hairy roots, every replicate was considered since that data is noisier). For plot ‘a’ (‘Human’), all normalized deletion\insertion frequencies that were used to create plots ‘e’-‘h’ (‘K562’, ‘T’, ‘U937’, ‘PLX’) were added together and normalized once more (divided by 4).

Plot ‘i’ was made using data from long sequencing that consequently showed much more deletions of longer lengths compared to data from short sequencing. In addition, some of the experiments had 2 sites for 2 different gRNAs in the same amplicon (Fig. S1A). Amplicons’ lengths in the double-guide experiments were usually larger compared to the single-guide experiments (Fig. S1B). This caused an artificial increase (bias) in deletion lengths of the size of the distances between the 2 sites when looking at the raw unfiltered data (Fig. S1C). To account for this, the data from experiments that had a single gRNA (‘SG’ - 18/103) were used to estimate the likely maximal length of deletions (Fig. S1C) in the double-guides experiments (‘DG’). Only deletions in the top 99% of the data for the SG experiments (i.e. deletions up to 504 nts) were extracted from the data of the DG experiments. Therefore, while longer deletions in the DG experiments that were not present in the SG experiments were not considered for the plot (which perhaps caused us to miss longer deletions) there was still a biased increase in longer deletion lengths due to A. smaller deletions overlapping in DG experiments and read as a single large deletion; B. Large deletions due to two cuts in the genome that caused the entire sequence between them to be deleted.

Despite these two biases, we still observed a signal for a negative monotonous relationship between frequency and deletion lengths, thus strengthening this claim about the data. The insertions histograms seemed relatively similar between both single and double gRNAs experiments (∼99% of insertions were up to 13 base-pairs in the single site experiments compared to ∼97% if when looking at both single and double sites) so no further processing was done on insertions.

Since the frequencies in the deletion lengths were vastly different in the different sequencing techniques (in an order of magnitude Fig. 7C (h-i)), combining the data to show some image of the data for tomato as a whole would have created a distorted picture, since longer reads than ∼20 are very rarely read in the short sequencing data.

## 2. Feature generation

### 2.1 Thermodynamics and structure features

Enthalpy-based features were calculated, under the assumption that they indicate the folding strength of the target site. Given a sequence, each two adjacent nucleotides are given a score representing the thermodynamics between them (the target’s thermodynamics). The features were: “enthalpy_mean” - the mean enthalpy calculated over all 23 nucleotides in target; “enthalpy_mean_ext” - the mean enthalpy calculated over the extended sequence (100 upstream, 100 downstream of target, total of 223 nt); “enthalpy_mean_up\down_ext” - the mean enthalpy calculated over the 100 nts up\downstream of the site; “enthalpy_x” - according to position of dimer, positions 1-22 in the target site.

Local DNA shape features were calculated using DNAshape (18) and designed to predict DNA shape attributes from 5 nucleotides: Minimal Grove Width (MGW), Helix Twist (HelT), Propellor Twist (ProT), and Roll as defined by Curves (52). For each position in the target site, we calculated these attributes based on the 5 nucleotides centered around that position. The actual features were similarly calculated as before: “DNAshape_{ATTRIBUTE}_mean” - mean score of ATTRIBUTE over all possible pentamers (19) in the site; “DNAshape_{ATTRIBUTE}_mean_ext” - the mean ATTRIBUTE over the pentamers calculated over the extended sequence (100 upstream, 100 downstream of target, total of 223 nt); “DNAshape_{ATTRIBUTE}_mean_up\down_ext” - the mean ATTRIBUTE calculated over the pentamers 100 nts up\downstream of the site; “DNAshape_{ATTRIBUTE}_x” - according to position of center of the pentamer, positions 1-19 in the target site.

RNAfold from ViennaRNA (19, 20) was used to check if the 1^st^\2^nd^\3^rd^\all the stems in the scaffold (given the spacer) (“Head1\2\3”, “1&2&3”). We also calculated the free energy of the gRNA (“guideEne”) and the free energy of the guide+scaffold (“guide&scafEne”). In addition we had a binary feature depicting if the nucleotides of gRNA are paired to the nucleotides of the scaffold (“isBasePairs”). We also calculated the free energy of the site + 10 nts upstream and downstream based on MATLAB’s oligoprop function by averaging 3^rd^ column in the Thermo property of the results (“DNA_fold”).

### 2.2 Expression features

We wanted to consider the expression of the genes in which the editing sites were in. Since we didn’t have these measurements for all organisms, we used CAI (21) and ChimeraARS score (22), as well as simple codon and amino acid frequency, as a proxy. For each site, a symmetric window of win_$w nts on each side (w = 0, 10, 20) was taken and cross-referenced with all coding genes. If at least one gene’s ORF intersects with the window, all features are calculated on the overlapping codons.

A site may intersect multiple genes with different reading frames; thus, for features 1-4 we calculate the (a) average score of all genes overlapping with the site; (b) maximal such score;

- .aa_freqs For each codon we first calculate the genomic frequency of its corresponding amino acid, i.e. its occurrences in coding genes relative to other AAs. aa_freqs is the geometric mean, for the overlapping codons, of this frequency.
- .codon_freqs Similar to aa_freqs, using the genomic frequency of the codon itself instead of the AA’s frequency.
- .CAI Similar to features 1-2; the CAI (codon adaptation index) (21) is the frequency of the codon in some reference set, normalized to the frequency of the most abundant synonymous codon. We chose the whole genome to be the reference set.

The chimeraARS score (22), based on the entire genome. Briefly, the chimeraARS score assesses compatibility between a sequence (our window) and a reference set (highly expressed genes) by finding the average length of their longest common substrings.

### 2.3 Epigenetics and accessibility features

The efficiency of CRISPR/Cas9 is influenced by several epigenetic factors, including chromatin accessibility (using DNase-seq), CTCF binding, H3K4me3, and DNA methylation, and was previously modeled (9, 53). The features based on these epigenetic factors were created by checking the overlap between the site and known CTCF-defined boundaries, sites where the DNA was found to be methylated from Reduced Representation Bisulfite Sequencing (RRBS) data, sites where histone H3 was trimethylated at lysine 4, and hyper-sensitive DNase sites. These features were only available for humans.

Chromatin accessibility is the degree to which nuclear macromolecules can physically contact chromatinized DNA and is determined by the occupancy and topological organization of nucleosomes as well as other chromatin-binding factors that occlude access to DNA (54). Open or accessible regions of the genome are, thus, regarded as the primary positions for regulatory elements (55). Hence, the accessibility of the target DNA sequence in the genome plays a crucial role in the success of Cas9-mediated cleavage and subsequent editing. gRNA that targets genome loci at an accessible region of the genome, can better access the target site and form a stable complex with Cas9, leading to improved cut efficiencies.

CTCF is a ubiquitously expressed and highly conserved protein involved in chromatin remodeling and organization. One of the important roles of CTCF is binding at chromatin domain boundaries and defining insulator regions (56).

These insulator regions can maintain an open chromatin conformation around genes. If the target site is located within a CTCF-defined boundary, it is more likely to result in improved cut efficiencies. Moreover, CTCF binding can lead to remodeling of the chromatin and to open chromatin which is generally more favorable for Cas9 cutting.

Trimethylation of Histone H3 at Lysine 4 (H3K4me3) is a major chromatin modification associated with open and accessible chromatin regions (57). In open chromatin, the target DNA is more accessible to the CRISPR-Cas9 complex, making it easier for Cas9 to bind and create a double-strand break (DSB). Consequently, this modification contributes to increased cut efficiencies.

DNA methylation is an epigenetic mechanism that involves the addition of a methyl group to cytosine residues in the DNA sequence, which plays a critical role in gene regulation and chromatin structure (58). The presence of methyl groups on cytosine residues can create steric hindrance, making it challenging for Cas9 to access and bind the target sequence. Hence, it physically hinders the binding of Cas9 to the target site, reducing the cut efficiency in methylated regions. Locations in the genome that were methylated were measured using Reduced Representation Bisulfite Sequencing (RRBS) sequencing.

DNase I (deoxyribonuclease I) is an endonuclease that cleaves DNA nonspecifically and is commonly used in genomic assays to probe chromatin accessibility. It preferentially digests regions of open chromatin, where DNA is less tightly bound by nucleosomes and other chromatin-associated proteins, allowing for greater enzyme access. Using Dnase-seq data, we found hypersensitive sites to Dnase that are believed to be in regions of open chromatin.

Another set of features was generated using TADbit (59) to map raw Hi-C reads, create an interaction matrix and create a 3D model of the chromosomes. Each chromosome is divided into particles with 100kb resolution, i.e. particle #1 is coordinates 1-100, 000; particle #2 is 100, 001-200, 000; et cetera. The site’s score is the score of its corresponding particle. More information about this model can be found here (16). The features were “tad_density” - Chromatin density (number of nucleotides per chromatin nm); “tad_angle” - Angle between 3 adjacent particles; “tad_interactions” - Number of particle interactions (number of particles closer than a certain cutoff).

For the methylation features, we also utilized the methylation motifs found in (23). These motifs describe the probability of a sequence to match an empirical methylation site. Thus, we used a sliding window method and compared each window with all motifs, and the highest score found was reported. We expected these values to inversely correlate with editing efficiency – sites that are more likely to be methylated, have a higher likelihood to be folded, and thus inaccessible to gRNAs.

### 2.4 Metagenomic features

This set of features compared the signatures found in gRNAs to the ones found in naturally occurring CRISPR arrays in microorganisms. To that end, the assemblies of all publicly available genomes and metagenomes, excluding genomes of Fungi, Metazoa, and Viridiplantae, were downloaded from NCBI WGS (60) and EBI Mgnify (61) on March 14, 2020. Overall, the dataset included 596, 338 genomes and 22, 923 metagenomes. CRISPR arrays were detected using MinCED (62), and the spacers were de-duplicated. The Cas genes associated with the arrays were identified using a set of hidden Markov models describing Cas protein families (63). From these data five sets of spacers were compiled to computer the metagenomic signatures: (i) full spacer set; (ii) spacers associated with Cas9; (iii) spacer associated with type II-A Cas9; (iv) spacers associated with Cas9 protein that shares at least 50% identity with Cas9 from Streptococcus pyogenes (SpyCas9) (64); and (iv) spacers associated with Cas9 protein that shares at least 90% identity with SpyCas9. Next, the distribution of nucleotide 3-mers and 5-mers of nucleotide in these five sets was computed, i.e., the frequency of each 3-mer and 5-mer in each spacer set. According to their frequency, the k-mers were then assigned a label according to their quartile (Q1, Q2, Q3, or Q4). For a given gRNA, the abundance of its k-mers in nature was quantified according to their frequency in each of the quartiles pre-calculated from the spacers we found in metagenomics. This yielded 40 features per gRNA: the frequency of each one of the 4 quartiles x 5 sets of spacers x 2 sizes of k-mer (3-mers and 5-mers). In addition, we also calculate the log of the probability to observe the set of the gRNA k-mers in each of the natural sets of spacers.

This was achieved by summing the log frequencies of each of the k-mers of the gRNA. This yielded 10 additional features (5 sets of spacers x 2 sized of k-mers). The abovementioned features were also calculated just a reduced alphabet representing GC vs AT, i.e., all Gs and Cs were represented by one character, and all As and Ts by a second character. The features described above were calculated on this reduced alphabet representing strong vs. weak hydrogen bonds.

### 2.5 Sequence features

Local Composition Complexity (LCC) is a local metric, defined over a sliding window. It is defined as the KL divergence between the nucleotide count and a uniform distribution of nucleotides (24). This feature has a highly optimized implementation in the Biopython library. We expected these features to correlate with editing efficiency windows that have high LCC values have a more distinct, unique signal, that might be more easily recognized by the gRNA.

### 2.6 Competition features

Competition-based features capture the potential influence of other genomic sites on Cas9 activity at a given target. These features quantify the number of sequences in the genome that are like the target site, which could compete for binding by the same gRNA. If numerous similar sites are distributed throughout the genome, particularly at greater distances from the target, they may reduce Cas9 binding efficiency at the intended site by diverting binding events elsewhere, especially when that off-targets are next to a PAM-motif (65). Conversely, when such competing sites are in proximity of the target site, i.e. within approximately 1, 000 nucleotides, they may help recruit Cas9 to the local genomic region, increasing the likelihood of cleavage at the target. The features are:

A. Number of sites matching the PAM-adjacent target site suffix of suffix_length (5, 10, 15, 20) that are followed by an NGG-PAM motif (if PAM exists – the name in feature contains ‘ngg_pam’, otherwise it does not), allowing for a Levenshtein distance of up to d (0, 1, 2, 3, 5). ref_seq is the search space (genome, ORF, full gene - including introns). The name of the feature in the figures\tables is sites_count_ngg_pam_{ref_seq}_-{suffix_length}_{d} where ref_seq\suffix_length\d can be any of the values described here.
B. Number of local sites matching the PAM-adjacent target site suffix of suffix_length (0-PAM only, 1, 2, 3, 5) that are followed by an NGG-PAM motif or not (if exists name in feature contains ngg_pam, otherwise nothing), within a maximum distance of max_distance (0, 10, 100, 1000, 10000, 100000, 1000000, 2, 25, 250, 2500, 25000, 250000) to the target site. The name of the feature in the figures\tables is local_sites_count_{ngg_pam}_{max_distance}_-{suffix_length} - where ngg_pam\max_distance\suffix_length can be any of the values described here.

## 3. Statistical tests used for analyses

### 3.1 Combination of models analysis

For each dataset separately, we trained Linear, XGBoost, and Random Forest models on randomly selected 80% of the data and tested on the remaining 20%. This procedure was repeated 100 times with every possible combination of the predictions from the 4 public models (CRISPRedict, SPROUT, DeepCRISPR, and uCRISPR) as features. The type of best chosen model using a single feature (in light blue) is compared to the best chosen model using every combination of features (in dark blue). The p-value was calculated using Wilcoxon signed-rank test and were adjusted for multiple tests using the Benjamini-Hochberg method.

The data for uCRISPR used in this analysis is a subset of the data used to train uCRISPR and came from functional screening in A375 cells. The data for SPROUT used used in this analysis is the data used to train SPROUT and came from endogenous editing in T cells. The data for DeepCRISPR used used in this analysis is part of the data used to train DeepCRISPR and came from functional screening in HEK293 cells. The data for CRISPRedict used in this analysis is the data used to train CRISPRedict and came from functional screening in HEK293 cells.

### 3.2 Partial correlation analysis

To obtain robust significance estimates that accommodate dependencies among features, we computed empirical p-values from repeated permutations of the editing-efficiency vector. For each of 10, 000 permutations, we recalculated all partial correlations and quantified, for each observed correlation, the proportion of randomized values with equal or more extreme magnitude. Significant results are summarized in Table S11. To further control for false discoveries, we estimated an empirical FDR by repeating this procedure 100 times, each time selecting a random permutation-derived partial Spearman correlation vector, computing empirical p-values, and counting how many features in that cell type would be deemed significant at a threshold of 0.01. We then averaged the number of significant features across permutations and divided it by the number of significant features observed in the real data, yielding an empirical estimate of the expected FDR for each system (Table S12).

### 3.3 System-specific determinants of Cas9 efficiency analysis

To compute empirical p-values, we repeated the full analysis 1, 316 times with permuted editing-efficiency labels (chosen to avoid empirical p-values of zero). For each feature and direction, we calculated the empirical p-value as the fraction of permutations whose one-sided Mann–Whitney p-value was less than or equal to the corresponding real-data p-value. Using the permutation distribution, we estimated the FDR and selected an empirical significance cutoff that ensured an expected FDR <10% in all cell types.

### 3.4 Repair profiles are universal in various cellular systems statistical analysis

To support our claim that there are more deletions than insertions, we performed a binomial test seeing how likely it is to see the number of successes (or more) that were defined as the fraction of reads that were deletions out of 10000, where n was set to 10000. This is a very lenient assumption as the number of reads in the short sequencing is orders of magnitude higher, but in the long-sequencing data i.e. tomato hairy roots, that is the order of magnitude for the number of reads. The chance to see a deletion according to this null model was 0.5.

To support our claim that there are more short deletions than long deletions, we calculated the Spearman correlation of the deletion lengths with their frequencies. The correlations and the p-values: Fly r= -0.97, p= 5e-72; K562 r= -0.81, p= 2.9e-38; T r= -0.98, p= 1.1e-90; U937 r= -0.90, p= 1.4e-51; Prawn r= -0.77, p= 2.0e-24; PLX r= -0.91, p= 3.2e-47; Tomato leaf r= -0.61, p= 2.5e-12; Human r= -0.95, p= 9.0e-80; Tomato hairy roots r= -0.86, p= 1.4e-164.

To support our claim that there is a clear advantage for 1-bp insertions, we performed a binomal test between the number of 1-bp insertions (n_bp_1= sum of percentages of reads times 10000 - number of reads) and the number of every other insertion length (n_bp_x), where the probability of success was 0.5, number of trials was n_bp_1 + n_bp_x, and number of successes was n_bp_1. For each cell we took the maximal p-value (after BH correction). We then performed BH correction on the p-values for each cell. The p-values for the claim that 1-bp insertions are more frequent than any other length of insertion is: Fly p=3e-09; K562 p=2e-230; T p=6e-190; U937 p=4e-82; Prawn p=1e-69; PLX p=1e-259; Tomato leaf p=1e-259; Human p=1e-225; Tomato hairy roots p=1e-122.

### 3.5 The identity of the 1-bp insertions is usually a copy of the nt upstream to the cut-site analysis

To test whether 1-bp insertions preferentially copy the nucleotide immediately upstream of the Cas9 cut site, we performed a localized permutation test that controls for the sequence composition around each editing event. For every 1-bp insertion event, we first recorded the true upstream base and the identity of the inserted base, and we also extracted a ±5 bp window around the cut site from the reference sequence. The observed dataset was summarized as a 4×4 matrix giving, for each upstream nucleotide, the normalized frequency of each inserted nucleotide. To generate a null distribution that preserves the local sequence context, we created randomized datasets by replacing the true upstream base of each event with a nucleotide sampled uniformly at random from its local ±5 bp window (excluding the cut position). This procedure was repeated many times (1, 000 permutations), yielding a distribution of expected insertion patterns under a model where insertion identity reflects only the local nucleotide composition and not specifically the upstream base. The observed upstream-matching rate was then compared to this null distribution to compute an empirical p-value, quantifying whether 1-bp insertions copy the true upstream nucleotide more often than expected by chance given the local sequence context. All p-values were less than 0.001 after correcting for FDR.

## 4. Experimental methods

All data other than the hairy roots data were amplified by PCR using primers with Illumina-compatible overhang adapters, Samples were prepared for Illumina MiSeq (Illumina, San Diego, CA, USA) according to the manufacturer’s instructions and sequenced using 2 x 150 bp paired-end reads.

### 4.1 T cells

#### 4.1.1 Knockout of genes in T cells

T cells were isolated using MACS beads and activated for 48 hrs. 0.5 * 10^6^ cells were washed in PBS and resuspended in20 µl of P3 buffer for electroporation (Lonza). RNA-protein particles (RNPs) were assembled by mixing 1.3 µl of Cas9 (IDT) with 160 pmole sgRNA (IDT) and PBS to a final volume of 6 µl. the mixture was incubated for 10 min at 25°C and kept on ice until use. Prior to electroporation 1 µl of 100 µM Enhancer (IDT) was added to the mixture and the mixture was added on top of the cells. The cell-RNP-Enhancer mixture was transferred to an electroporation cuvette.

Electroporation was performed in Lonza 4D nucleofector using the EO-115 program. Electroporated cells were allowed to recover in T cell complete medium containing 200u/ml IL-2. Cells were harvested after 72hrs of recovery and crude DNA extract were prepared and subjected to library preparation.

### 4.2 K562

#### 4.2.1 K562 CRISPR/Cas9 Genome Editing

The K562 cell line was obtained from ATCC and cultured in RPMI medium supplemented with 10% FBS. CRISPR/Cas9 gene editing was carried out using the Alt-R® CRISPR-Cas9 enzyme (IDT), sgRNA guides (IDT), designed with the “GoGenome” software developed by the CRISPRIL consortium and the electroporation enhancer (IDT). Briefly: CRISPR Mix of the Cas9 enzyme (120 pmol) was combined with 250 pmol of sgRNA guide and incubated for a 20-minute. Electroporation enhancer (1 nmol) was added immediately before electroporation. After washing the cells with PBS, they were resuspended in 20 µL of electroporation buffer SF (LONZA) mixed with the CRISPR components. Electroporation was performed using the Lonza-4D electroporation program (FF-120). After electroporation cells cultured in RPMI medium supplemented with 10% FBS.

#### 4.2.2 Post-Electroporation Analysis

Forty-eight hours following electroporation, cells were harvested and DNA was extracted to evaluate editing efficiency and variations. Guide editing was assessed by next-generation sequencing (NGS) through a two-step PCR amplification process using the 18S Illumina Amplicon Protocol. The first PCR used specific primers, followed by a second PCR step to add barcodes. The PCR product size and concentration were analyzed using a D1000 TapeStation and Qubit. NGS library preparation was performed at 8 pM concentration with 20% PhiX, and sequencing was conducted on a MiSeq platform with a 2x151 read configuration.

### 4.3 U937

U937 cells (a kind gift from Prof Ofer Medelboim) were transduced with lentiCas9-Blast (Addgene #52962) lentivirus and selected with 40ug/ml of Blasticidine. U937-Cas9 cells were transduced with single guide lentivirus and selected using Puromycin (4ug/ml). The selected U937-Cas9 cells were differentiated into macropahges by PMA (100ng/ml) for 5hrs, then washed and further differentiated for 48hrs into M2 macrophages using a combination of cytokines IL10/IL4/TGFb (10ng/ml each from PeproTech), after 48hrs we extract genomic DNA using GeneAll DNA extraction kit (GeneAll Biotechnology Co., Ltd, Korea). Genomic DNA sequences flanking the cut sites were amplified by PCR using primers with Illumina-compatible overhang adapters, Samples were prepared for Illumina MiSeq (Illumina, San Diego, CA, USA) according to the manufacturer’s instructions and sequenced using 2 x 150 bp paired-end reads.

### 4.4 PLX

#### 4.4.1 Placental Cell Preparation

Intermediate cell stocks (ICS) of maternal or fetal placenta-derived mesenchymal-like adherent stromal cells were produced by Pluri-biotech Ltd. (Haifa, Israel). In the production process, the cells are digested from donated placental tissue, expanded in large tissue culture flasks, and cryopreserved as intermediate cell stock, which is then transferred to bioreactors for further controlled 3D-expansion on non-woven polyester and polypropylene-made carriers.

#### 4.4.2 Generation of CRISPR-Cas9 Knockout cells

Ribonucleoprotein (RNP) complexes were created by mixing 10^5^ pmol of Alt-R Cas9 protein (IDT; Coralville, IA, USA) with 260 pmole of Alt-R sgRNAs (IDT; Coralville, IA, USA) in PBS, and incubation at 25°c for 15 minutes. Immediately before electroporation, 0.5 * 10^6^ 2D-expanded ICS of placental cells in 20µl of P1 buffer (Lonza, Basel, Switzerland) were combined with 6µl of RNP complexes, and transferred to Nucleocuvette strips (Lonza, Basel, Switzerland). Cells were nucleofected using pulse code CA-137 in a 4D-Nucleofector X unit (Lonza, Basel, Switzerland). Nucleofected cells were further expanded in tissue culture flasks, and finally harvested and cryopreserved.

#### 4.4.3 Evaluation of CRISPR-Cas9 editing efficiency

Prior to freezing, genomic DNA was extracted from 0.5x10^6^ cells using QuickExtract DNA extraction solution (Lucigen, Middleton, WI, USA). Genomic DNA sequences flanking the cut sites were amplified by PCR using primers with Illumina-compatible overhang adapters, Samples were prepared for Illumina MiSeq (Illumina, San Diego, CA, USA) according to the manufacturer’s instructions and sequenced using 2 x 150 bp paired-end reads.

### 4.5 Tomato hairy roots

#### 4.5.1 Plant material and generation of genome-edited hairy roots

Tomato seedlings (*Solanum lycopersicum* cv. M82) were grown in a walk-in tissue culture room at 24 °C under a 16/8 h light/dark photoperiod. Genome-edited hairy roots were generated following a previously reported protocol (66), with minor modifications. Briefly, tomato seeds were surface-sterilized in 70% ethanol for 3 min, followed by 3% (v/v) commercial bleach (original concentration 6%) for 15 min, and then rinsed six times with sterile distilled water. Sterilized seeds were germinated on half-strength MS medium (4.5 g L^-1^ MS222, Ducheffa, The Netherlands; 1.5% sucrose; 0.8% plant agar; pH 5.8) in Magenta boxes.

After 7–10 days of growth, cotyledons were excised and placed on co-cultivation medium (4.5 g L^-1^ MS222, 3% sucrose, 0.8% plant agar, and 200 µM acetosyringone; pH 5.8) and incubated in the dark for 1 day. Cotyledons were then immersed for 15 min with gentle shaking in a suspension of Agrobacterium rhizogenes strain ATCC 15834 harboring the genome-editing constructs, prepared in liquid MS222 medium (4.5 g L^-1^ MS222, 3% sucrose, and 200 µM acetosyringone; pH 5.8). Following infiltration, cotyledons were transferred onto co-cultivation medium plates overlaid with a sterile filter paper and incubated in the dark for 2 days. Subsequently, cotyledons were transferred to selection medium (4.5 g L^-1^ MS222, 3% sucrose, 0.8% plant agar; pH 5.8) supplemented with 250 mg L^-1^ cefotaxime and 50 mg L^-1^ kanamycin, and grown in the walk-in tissue culture room. Hairy roots typically emerged from the cut surfaces of the cotyledons within 10 days.

#### 4.5.2. Generation of genome-editing constructs

Genome-editing constructs were generated as previously described (67). Using a two-step PCR-based strategy, individual sgRNAs were cloned into expression cassettes driven by the Arabidopsis thaliana U3d, U3b, U6-1, or U6-29 promoters in an intermediate plasmid. The resulting sgRNA expression cassettes were subsequently assembled into the w1-35Spro-GFP-Cas9-CCD binary vector using Golden Gate ligation cloning.

#### 4.5.3. Multiplex PCR-based NGS assay

On-target genome-editing efficiency was assessed using multiplex amplicon sequencing. SMRTbell libraries were prepared using the PacBio Barcoded Overhang Adapters Kit 8A (101-628-400, PacBio, USA) for long-read sequencing. Primers were designed to amplify genomic regions spanning approximately 500 bp upstream and downstream of each, or both, sgRNA target sites. Genomic DNA was extracted from hairy roots using the cetyltrimethylammonium bromide (CTAB) method as previously described (68).

PCR amplification was performed using Q5® High-Fidelity DNA Polymerase (M0492S, NEB, USA) with sgRNA-flanking primers and genomic DNA as template. PCR products were purified using AMPure PB Beads (100-265-900, PacBio, USA) to remove residual primers, enzymes, and nonspecific products. Purified amplicons were quantified and pooled at equimolar concentrations. For each target amplicon, six biological replicates derived from A. rhizogenes-mediated transformation with genome-editing constructs and one control replicate transformed with A. rhizogenes competent cells were included, yielding seven pools.

Pools were barcoded using the PacBio Barcoded Overhang Adapters Kit 8B (101-628-500, PacBio, USA) according to the manufacturer’s instructions. Barcodes 1-6 were replicates for mutated runs i.e. with CRISPR Cas9, while barcode 7 was used for WT (without CRISPR Cas9). Adapter ligation reactions (∼16.3 µL each) were combined into a single tube and purified with AMPure PB beads. Sequencing primers, Sequel® Polymerase 3.0, dNTPs, DTT, and binding buffer were added using the SMRTbell Express Template Prep Kit 2.0 (100-938-900, PacBio, USA), followed by incubation at 30 °C for 1 h to generate polymerase-bound SMRTbell® complexes. These complexes were purified with AMPure PB beads.

Internal control complexes were diluted sequentially (1:100), and sequencing samples were prepared by mixing 67.9 µL Sequel® Complex Dilution Buffer, 4.8 µL prepared sample, 2.8 µL diluted internal control complex, 8.5 µL DTT, and 1 µL Sequel® Additive. The final mixture (85 µL) was loaded per well for PacBio sequencing.

### 4.6 Rahan (Tomato leaf protoplasts)

gRNAs for both on-targets and off-targets and the primers used for amplification of region of interest, from transformed protoplast extracted genomic DNA, were designed using the GoGenome tool.

Expanded leaves of M82 tomato from 3.5-4 week-old seedling were cut into small pieces, submerged in 15 ml enzyme mixture (20 mM MES pH 5.7, 20 mM KCl, 10 mM CaCl2, 1.1 g Mannitol, 225 mg Cellulase R10 (Duchefa), 60 mg Macerozyme (Duchefa) and 15 mg BSA) and incubated in the dark for overnight with mild agitation (55 rpm). After verification of protoplast release under a microscope, digested leaves were filtered through a mesh strainer (100 µm). Strained protoplast mixture was transformed into a 13 ml tube and was centrifuged at 760 RPM for 10 min. After removal of most of the liquid, 3 ml of W5 solution (154 mM NaCl, 125 mM CaCl2, 5 mM KCl and 2 mM MES pH 5.7) was added to the protoplast pellet and protoplast were released by gently tapping at the bottom of the tube.

For protoplast purification, protoplast/enzymes/W5 mixture was added gently into a 13 ml tube on top of 5 ml sucrose solution (prepared by dissolving 11.5 g sucrose in final volume 50 ml RNase free water) and was centrifuged at 760 RPM for 10 min. The obtained protoplast phase (middle green band) was released into a new 13 ml tube, and was completed with addition of W5 solution up to the ∼ 12 ml mark. Followed by tube inversion, 20 µl of the resuspended protoplast were removed and count on a hemacytometer. The rest of the tube content was centrifuged at 760 RPM for 10 min. This was followed by resuspension of protoplasts pellet in MMG solution (0.4 M Mannitol, 15 mM MgCl2, 4 mM MES pH 5.7) to obtain cell density of 1 million protoplasts/ml.

To assemble ∼ 20 µg of sgRNA duplex, tracrRNA and crRNA (IDT) were dissolved in TE buffer (pH 8.0) to a concentration of 200 µM each, were mixed in equal molar concentration in the presence of nuclease-free duplex buffer (IDT) and incubated at 95°C for 5 min followed by room temperature for 10 min. To assemble the RNP, sgRNA duplex assembly mixture, was mixed with 10 µg of Cas9 protein (Sigma-Aldrich) in the presence of 1 X NEB-buffer 3.1 and incubated at room temperature for 15 min.

For PEG-mediated transformation, 200 µl of protoplasts in MMG solution (200, 000 protoplasts) were mixed with 20 µl of RNP complex solution and was added with 220 µl freshly prepared PEG solution (40% [W/V] PEG4000 (Sigma-Aldrich), 0.2 M Mannitol and 0.1 M CaCl2). For preparation of negative control samples, 20 µl of RNase free water were add instead of the RNP complex solution. The transformation mixture was incubated in the dark for 20 min at 25°C, followed by addition of 950 µl W5 solution and incubation at room temperature for 15 min. This was followed by centrifugation at 450xg for 5 min and removal of the upper liquid. Finally, protoplasts were resuspended in 1 ml WI solution (0.5 M Mannitol, 20 mM KCl and 4 mM MES pH 5.7) and were kept in the dark for 48 h.

After 48 h, transformed protoplast were centrifuged at 450xg for 5 min following removal of the upper liquid genomic DNA was extracted (plant, Geneaid). Genomic region of interest was amplified using Platinum SuperFiII green PCR master mix (Invitrogen) under the following conditions: 98°C/3min + 37 X (98°C/10 sec + 60°C/10 sec + 72°C/30 sec) + 72°C/min. Amplified products were sent for NGS sequencing at the Technion facility.

### 4.7 BetterSeeds (Tomato leaf protoplasts)

#### 4.7.1 Protoplast isolation and purification

Tomato seeds were surface-sterilized and sown on solidified media. First true leaves of 14 to 20-day-old seedlings were sliced into thin strips and immersed in 15 ml of enzyme solution (3.65 g Mannitol, 2 ml 0.5 M MES pH 5.7, 500 ul 2 M KCl, 0.75 g Cellulase “Onozuka” R-10 (Yakult Pharmaceutical), 0.2 g Macerozyme R-10 (Yakult Pharmaceutical), 500 ul 1 M CaCl2, 0.05 g BSA) in a Petri dish, and incubated in the dark overnight with mild shaking (25 RPM) for 16 h. Extracted protoplasts strained through a 0.1 um filter and washed with W5 solution (154 mM NaCl, 125 mM CaCl2, 5 mM KCl, 2 mM MES pH 5.7). Healthy, intact protoplasts were isolated using a sucrose gradient (23%, 11.5 g per 50 ml). After additional washing with W5, protoplasts were diluted to a concentration of 1 million cells per milliliter in MMG solution (0.4 M Mannitol, 15 mM MgCl2, 4 mM MES at pH 5.7).

#### 4.7.2 Assembly of CRISPR RNP

sgRNAs are purchased from Synthego as tracrRNA and target-specific crRNA. Both were diluted in TE buffer to a concentration of 200 uM each. Equimolar ratio of the two components were mixed with Duplexing Buffer and heated at 95oC for 5 min before slowly cooling to room temperature and stored at -20 C until day of transformation. SpCas9 protein was expressed from pET28b-Cas9-His68 (Addgene plasmid # 47327) by the Proteomics Unit at the Weizmann Institute. For each sample 10 ug of SpCas9 and 20 ug of duplexed sgRNA were mixed with NEB Buffer 3.1 at 20 ul total volume and incubated at room temperature for 15 min. RNPs were prepared immediately prior to transformation.

#### 4.7.3 PEG-mediated transformation of CRISPR RNP

20 ul preassembled RNP were added to 200, 000 protoplasts in 200 ul MMG solution. An equal volume of PEG solution (40% [w/v] PEG 4000 Sigma, 0.2 M Mannitol, 0.1 M CaCl2(was added and mixed gently prior to 20 min incubation in the dark. Then the samples were diluted with W5 solution and incubated for 15 min in the dark. The samples were centrifuged at 450xg with soft start/stop and resuspended in 1 ml WI solution (0.5 M Mannitol, 20 mM KCl, 4 mM MES pH 5.7) before incubation in the dark for 48 h.

#### 4.7.4 Sample collection and DNA extraction

Samples were pelleted using centrifugation at 450 x g. The WI was removed, and DNA was extracted using a crud plant DNA preparation procedure: Protoplast samples were incubated in 75µl DNA extraction buffer (100 mM NaOH, 2% Tween 20) for 10 min at 95oC and span-down. 75µl of second buffer (100mM Tris-HCl-pH 7.5, 2mM EDTA-pH 8.0) was added to the sample, the tubes were vortexed and span down. PCR was then performed using specific primers for the sgRNA target, together with Illumina Nextera Transposase Adapters: Nextera Read 1 Adapter (TCGTCGGCAGCGTCAGATGTGTATAAGAGACAG) and Nextera Read 2 Adapter (GTCTCGTGGGCTCGGAGATGTGTATAAGAGACAG).

### 4.8 Hazera (Tomato leaf protoplasts)

The protocol for the Tomato leaf protoplasts data acquired from Hazera is described in detail in the methods section of Ben-Tov et al (69).

### 4.9 Black Soldier Fly

The microinjection procedure was performed as described in (70). with necessary modifications. The injection mixture, composed of injection buffer, Cas9 protein (IDT), and gRNA (IDT), was incubated at 37 °C for 30 minutes prior to injection. Twenty freshly laid embryos were aligned on a slide and injected under a stereomicroscope using FemtoJet microinjector (Eppendorf). The injected embryos were then incubated at 30 °C and 90% relative humidity for two days. Subsequently, DNA was extracted from 5–10 embryos using the Quick-DNA Microprep Kit (Zymo Research), followed by PCR amplification of a 300 bp target region. A second PCR was performed to add sequencing adapters for library preparation. The libraries were sequenced using the Illumina MiSeq platform.

### 4.10 Prawn

The protocol for the data acquired from *Macrobrachium rosenbergii* primary cell culture (via electroporation), is the same as in Molcho et al (71).

#### 4.10.1 Animals and cells

All early developmental stage M. rosenbergii embryos used for injection and cell culture were acquired through the BGU breeding program, and a number of egg-carrying gravid females were obtained from Colors Farm Ltd. (Israel). These gravid females were produced in a tank system as follows: A single blue claw male was housed with four to six mature females in a 500-L tank. Each tank was checked daily for the typical male reproductive guarding behavior (72) as an indication of gravid females in the tank. Gravid females were then moved to a separate holding aquarium, and the embryos were examined under a light microscope to determine their stage, as described below. One- to four-cell embryos were used for the whole-animal editing platform, while embryos that had already passed the four-cell threshold were kept for 9–13 days up to the eyed-egg embryo stage and then used for the primary cell culture editing platform.

#### 4.10.2 Embryo stage identification

Embryos were delicately removed from the female with a fine forceps and examined under a light microscope (Nikon H550s, Nikon, Japan). The number of cells was determined by visual inspection. When visual confirmation of cell division was not adequate, cell division was shown by injecting the embryos with Fast Green FCF dye (Allied Chemicals, Thailand), which does not cross membranes, thereby enabling the visualization of cell divisions.

#### 4.10.3 Embryo preparation and sterilization

To prevent infection during incubation, fertilized eggs were removed from the brood chamber with fine forceps (or a toothbrush) and placed in a 50-mL tubes, each containing sterile salt water (15 ppt Coral Pro Salt; RedSea, Israel) and 10 μL of methylene blue. Following washing for 10 min in this solution, the eggs were rinsed with sterile salt water containing iodine (0.001 vol%) for 1 min, with sterile salt water containing formaldehyde (4%) for 5 min, and finally with sterile salt water for 5 min. The fertilized eggs were held in a small petri dish of salt water until injected.

#### 4.10.4 Embryo injections

A ribonucleoprotein (RNP)-Cas9 complex was prepared by mixing 9.5 pmol of synthesized single-guide RNA (sgRNA; IDT, USA) with 5.6 pmol of Cas9 protein (IDT) in a total volume of 1.8 μL. The complex was incubated for 5 min at room temperature before mixing with 0.05% phenol red to a total volume of 4 μL. For Cas9 mRNA delivery, 1 μg/μL GeneArt™ CRISPR Nuclease mRNA (A29378, ThermoFisher) was mixed with similar concentration of gRNA as mentioned above. All injections into M. rosenbergii fertilized eggs were performed with IVF micropipettes (Origio, USA), using a pneumatic PicoPump PV830, a vacuum injection system (WPI, USA), and two manual micromanipulators (WPI-M3301R/L, WPI). Pressure was set at 20 psi, and the injection duration set at 100 ms. By using a vacuum pump (Schwarzer Precision, Germany) at a negative pressure of 5 psi, eggs were kept in place for injection with a holding capillary. Then, the injection capillary was inserted into the egg, and successful insertion of the material was verified through the observation of a small phenol red stain at the moment of injection.

#### 4.10.5 Injection volume calculation

The PV830 injection system included a compressed hydrogen system to inject the RNP solution. The volume delivered per millisecond was calculated from the equations below using a known injection period of 500 ms followed by the measurement of the diameter of a drop of injected liquid on the tip of the injection capillary. The calculations below show that at an injection rate of 6.72 pL ms−1, 100 ms of injection would deliver 0.67 nL.

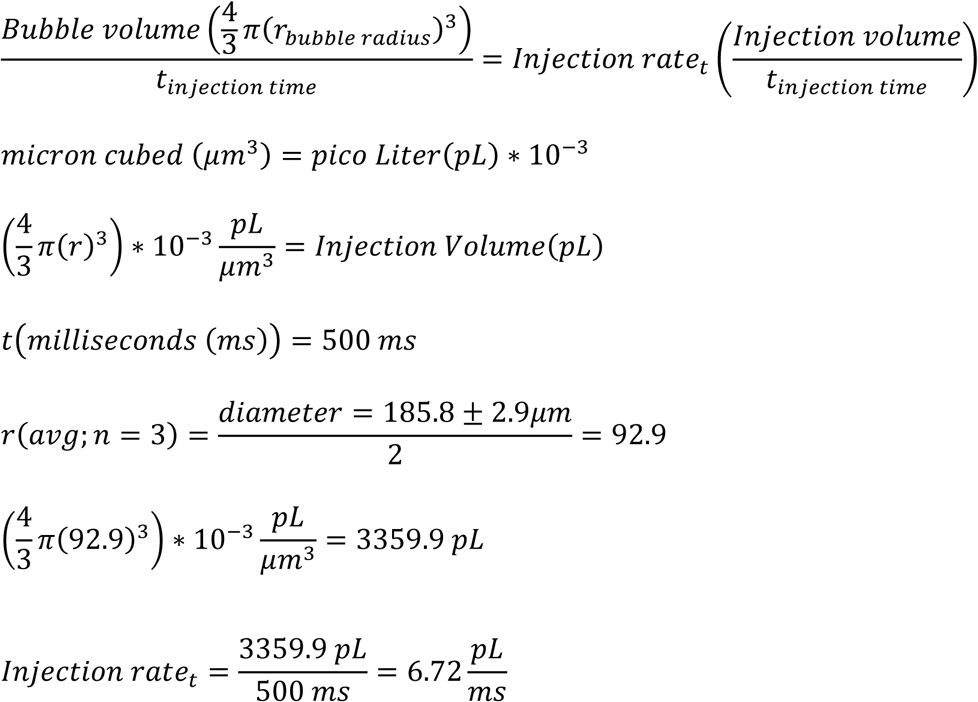

#### 4.10.6 Embryo care, DNA extraction, PCR amplification and amplicon sequencing

Injected embryos were transferred to a sterile incubator in small petri dishes, 20–30 embryos per dish, in an aqueous salt solution (15 ppt Coral Pro Salt) and kept at 27 °C for one week. The incubation solution was changed daily, and the eggs were examined visually for dead or malformed embryos. If infection (mainly fungal) was observed, non-contaminated eggs were moved to fresh dishes and further closely monitored. After one week, embryos were separated into individual tubes, and DNA was extracted as follows. Briefly, eggs were submerged in NaOH (0.2 N) and heated to 70 °C for 20 min in a Thermocycler Life Eco (Falc). After rough pipetting to ensure lysis, 50 μL of Tris HCl (0.04 M) were added to neutralize the acidity. DNA was then used as a template for amplification by PCR with specific primers flanking the gRNA. PCR products were cleaned using EPPiC Fast (A&A Biotechnology, Poland) and sent to the Macrogen sequencing lab (Amsterdam, Netherlands) for Sanger sequencing.

#### 4.10.7 Embryonic cell extraction, editing, and editing monitoring

Prior to the isolation of embryonic cells, M. rosenbergii egg-bearing females were disinfected in a methylene blue water bath (5 drops to 10 L water) at least one day prior to cell extraction. Eggs were further disinfected by washing for 10 min in crustacean physiological saline (CPS) (73) plus antibiotics (PEN/STREP, Biological Industries, Israel) and 0.5 μg/mL of the antifungal preparation Voriconazole (Sigma) in a rotator. Thereafter, the eggs were strained off in a 100-μm strainer (SPL Life Sciences, Republic of Korea) and washed for 1 min in 4% formalin-CPS solution, 1 min in 0.01% iodophor-CPS solution, and finally 10 min in CPS. Eggs were transferred to sterile Eppendorf tubes, containing 200 μL of Halt™ Protease and Phosphatase Inhibitor Cocktail (PI; Thermo-scientific) and 300 μL of CPS and homogenized. The homogenate was filtered through a 100-μm strainer, and the cells that passed through the strainer were collected in a 3-mL petri dish. Thereafter, these cells were transferred to a 15-mL tube and centrifuged at 850g for 6 min at 18 °C. The supernatant was discarded, and the pellet was resuspended in 1 mL of PI-CPS and then transferred to a Percoll step gradient comprising 3 mL of 100% physiological Percoll (13.5 mL commercial Percoll with 1.5 mL of CPS × 10), 2 mL of 50% Percoll (physiological Percoll diluted with CPS × 1), 4 mL of 25% Percoll, and 3 mL of 12.5% Percoll. Cells were centrifuged at 850g for 30 min at 18 °C. Cell fractions were transferred to a 15-mL tube and washed once with CPS. The cells were seeded at 3 * 10^6^ cells per well in a 6-well plate and grown in Opti-MEM Medium whose osmolality was adjusted to 420 mOsm with CPS. Plates were incubated at 28 °C with 5% CO2 for 24 h before RNP nucleofection. To monitor editing success, PCR products, using the genomic DNA as template, were sent to the Technion - Israel Institute of Technology (Israel) for next generation sequencing (Miseq Run V2; 2 × 150 bp, assuming 4 M reads per ends per run). The indels caused by the repair of the double-strand breaks through nonhomologous end-joining (NHEJ) were monitored.

#### 4.10.8 Cell nucleofection

Nucleofection was performed using an Amaxa P3 Primary Cell 16-well Nucleocuvette Strips kit (V4XP-3032; Lonza, Switzerland) and an Amaxa 4D-Nucleofecter (Lonza). The RNP Cas9 complex was formed by mixing 2.64 μL (163 pmol) of Cas9 62 μM (1, 081, 059; IDT) with 1.76 μL (176 pmol) of sgRNA 100 μM (IDT) in P3 nucleofection buffer (total volume of 11 μL); the mixture was incubated at room temperature for 20 min. For mRNA Cas9 delivery, GeneArt™ CRISPR Nuclease mRNA (A29378; ThermoFisher) was used in a final concentration of 0.06 μg/μL. An amount of 4 * 10^5^ primary cells were harvested, washed once in Opti-MEM, and resuspended in 28.6 μL of P3 nucleofection buffer. The cell suspension was mixed with RNP-Cas9, and 20 μL of the cell-RNP suspension was transferred to each of two wells of the 16-well strip nucleocuvette. Cells were electroporated using program CL-137 on the 4D-Nucleofector. Immediately after nucleofection, 80 μL of Opti-MEM + 10% FBS was added to each well at room temperature, and all 100 μL of cell suspension were transferred to a 96-well plate containing 50 μL of pre-warmed medium. The plate was incubated at 28 °C under 5% CO2. Edited cells were harvested three days after nucleofection. For DNA extraction, cells were suspended in culture medium and centrifuged at 11, 000g for 5 min. The supernatant was discarded, and the cells were resuspended in 10 μL of NaOH (0.2 N). Tubes were heated to 70 °C for 20 min and vortexed before the addition of 50 μL of Tris HCl (0.04 M).

## 5. Image generation for figures

All images in figure 1C were created with assistance from OpenAI’s DALL·E.

## 6. Data availability

All raw data can be accessed through http://www.ncbi.nlm.nih.gov/bioproject/1396917. All code and features for each cell can be viewed in https://github.com/shaicoh3n/CRISPR_maagad.

## Acknowledgements

We’d like to thank the Israeli Innovation Authority for funding this research through the CRISPRIL consortium. This study was supported in part by a fellowship from the Edmond J. Safra Center for Bioinformatics at Tel-Aviv University.

