## Supplementary information for "Context-dependent determinants of CRISPR-Cas9 editing efficiency revealed through cross-species endogenous editing analysis"

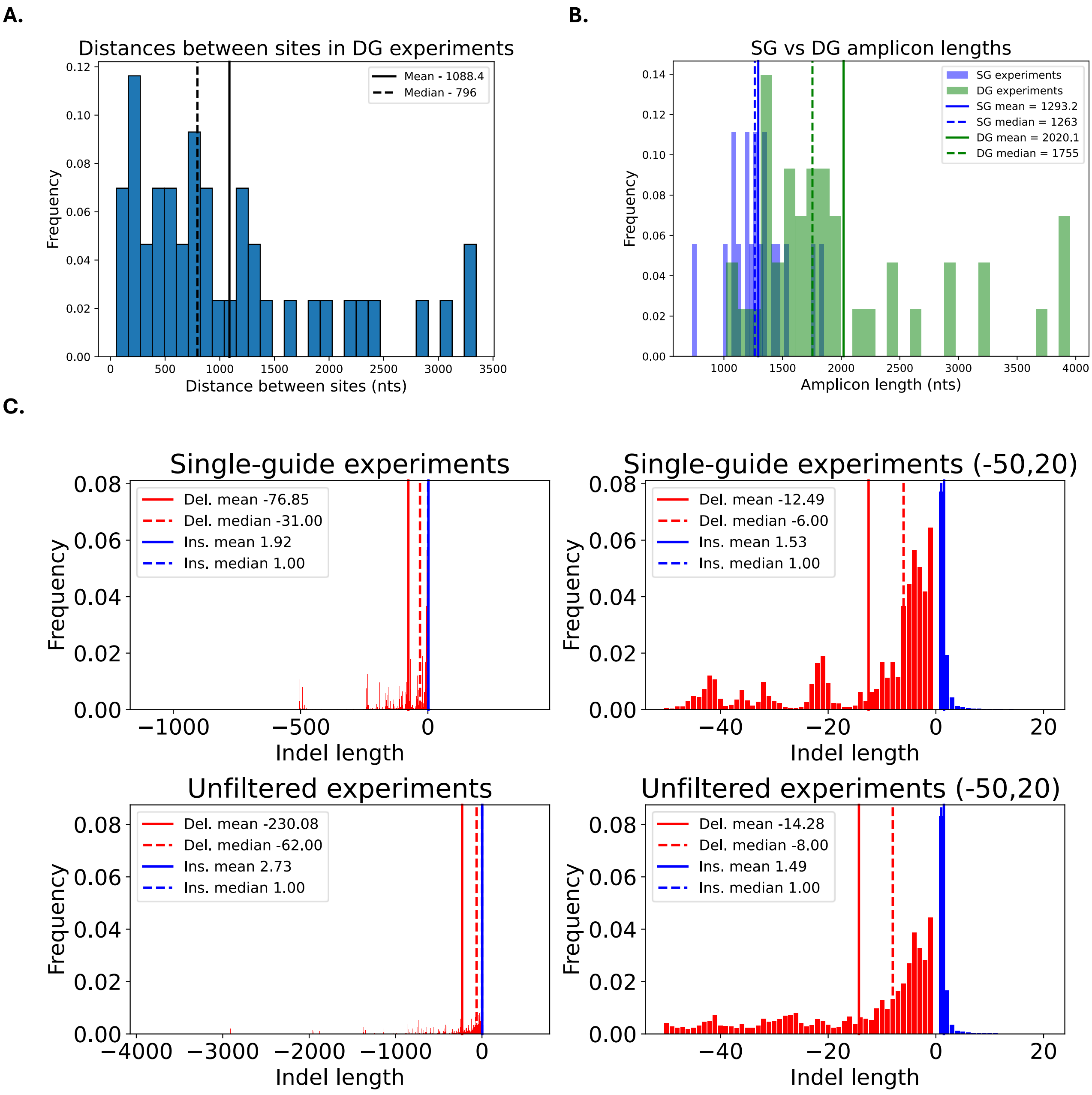


***Figure S1. Tomato hairy roots additional information.*** *'DG' stands for 'double guides' experiments and 'SG' stands for 'single guides' experiments. The product frequencies for all experiments of each cell type are summed according to the indel length. Positive indel lengths indicate insertions and negative lengths indicate deletions. The '(-50,20)' graphs are zoomed-in graph of the relevant h 'j' between deletions of length 1-50 and insertions between 1-20. 'Unfiltered experiments' graph contains all the raw data from the tomato hairy roots experiments (both DG and SG). The 'Single guide experiments' graph was made with the raw data from the single guides experiments and clearly shows that the vast majority (99%) of deletions are up to 504 nts.*
